# The NO Answer for Autism Spectrum Disorder

**DOI:** 10.1101/2023.01.07.523095

**Authors:** Manish Kumar Tripathi, Shashank Kumar Ojha, Maryam Kartawy, Wajeha Hamoudi, Adi Aran, Haitham Amal

**Affiliations:** Institute for Drug Research, School of Pharmacy, Faculty of Medicine, The Hebrew University of Jerusalem, Jerusalem, Israel; Neuropediatric Unit, Shaare Zedek Medical Center, Jerusalem, Israel; Faculty of Medicine, The Hebrew University of Jerusalem, Jerusalem, Israel

## Abstract

Autism spectrum disorders (ASDs) include a range of developmental disorders that share a core of neurobehavioral deficits manifested by abnormalities in social interactions, deficits in communication, restricted interests, and repetitive behaviors. Several reports showed that mutations in different high-risk ASD genes, including *SHANK3* and *CNTNAP2*, lead to ASD. However, to date, the underlying molecular mechanisms have not been deciphered, and no effective pharmacological treatment has been established for ASD. Recently, we reported a dramatic increase of nitric oxide (NO) in ASD mouse models. NO is a multifunctional neurotransmitter that plays a key role in different neurological disorders. However, its role in ASD has not yet been investigated. To reveal the novel molecular, cellular, and behavioral role of NO in ASD, we conducted multidisciplinary experiments using cellular and mouse models as well as clinical samples. First, we treated WT mice with an NO donor, which led to an autism-like phenotype. Next, we measured and found high levels of nitrosative stress biomarkers in both the *Shank3* and *Cntnap2* ASD mouse models. Treating both mouse models with a selective neuronal NO synthase (nNOS) inhibitor led to a reversal in the molecular, synaptic, and behavioral ASD phenotypes. Using a primary neuronal cell culture, we confirmed that NO is specifically involved in neurons in ASD pathology. Next, using genetic manipulations in the human SH-SY5Y cell line, we found that nNOS plays a key role in the pathology. Finally, we examined human plasma samples from 19 low-functioning ASD patients, compared to 20 typically developed volunteers, and found a significant elevation in the NO levels in the ASD patients. Furthermore, using the SNOTRAP technology, which is an innovative mass spectrometric method to identify the SNO-proteome (SNO: NO-mediated post-translational modification), we revealed that the complement systems in the synaptic and neuronal development processes are enriched in the ASD group. This work indicates, for the first time, that NO plays a pathological role in ASD development. Our findings will open future and novel directions to examine NO in diverse mutations on the autism spectrum as well as other neurodevelopmental disorders and psychiatric diseases. Most importantly, it suggests a novel treatment strategy for ASD.

**One sentence summary:** Nitric oxide plays a key role in ASD pathology development and progression, and targeting its production leads to a reversal in the autistic phenotype.

## Introduction

Autism spectrum disorder (ASD) is a neurodevelopmental and behavioral disorder (*1*); it is one of the most disabling conditions and chronic illnesses in childhood. It is characterized by abnormalities in social interactions, deficits in communication, restricted interests, and repetitive behavior (*2*). Once developed, this disorder descends to permanent lifelong abnormalities and disabilities. Globally, recent epidemiological studies referred to ASD prevalence rates of 1 in 44 children (*3*). The number of people with ASD is growing quickly. Thus, the prevalence rate has increased nearly three-fold over the last 20 years in the US alone. Unfortunately, to date, no effective pharmacological treatments or preventive measures are currently available for this spectrum disorder (*4*). ASD is a highly genetic pathology (*5*–*8*). Rare de novo mutations of genes such as *SHANK3*, *CNTNAP2*, and others appear to have functional effects on the protein-coding regions of the genome, representing a high risk of ASD (*9*, *10*).

Shank3 is a scaffolding protein located in the postsynaptic density complex of excitatory synapses (*11*). It binds to neuroligins and actin and participates in regulating actin polymerization, growth cone motility, dendritic spine morphology, and synaptic transmission (*12*). Deletions or mutations of the *SHANK3* gene have been found in patients with Phelan-McDermid Syndrome (PMS). Over 50% of these patients are diagnosed with ASD (*13*, *14*). This mutation is also found in ASD patients outside the PMS (*15*, *16*). Previous studies on *Shank3* mutant mice revealed ASD-like behaviors, such as impaired social interaction, anxiety-like behavior, and redued interest in novel objects (*17*, *18*). It is considered as one of the most promising mouse models of ASD to date (*16*, *19*, *20*).

The contactin-associated protein-like2 gene (*CNTNAP2*) encodes a neuronal transmembrane protein member of the neurexin superfamily, which is involved in neuron-glia interactions and clustering of K+ channels in myelinated axons. It plays a key role in synapse formation and stabilization (*21*, *22*). Moreover, it is considered one of the crucial genes in autism; it is responsible for delayed speech-language development and relatively impaired signaling (*23*, *24*). *Cntnap2* knockout mice display reduced stability in newly formed dendritic spines (*25*–*27*). Loss-of-function mutations in this gene have been implicated in ASD and cortical dysplasia–focal epilepsy syndrome (*28*). Studies on *Cntnap2* ^(*-/-*)^ mice revealed frequent spontaneous seizures and behavioral patterns typical of ASD, such as stereotypic motor movements, behavioral inflexibility, a reduced time of interaction with other mice in a Juvenile Playtest, and fewer isolation-induced ultrasonic vocalizations (distress calls) of the mouse pups to their mothers (*25*, *29*). The etiology of ASD remains elusive (*30*, *31*).

Recently, we found elevated levels of nitric oxide (NO) in an ASD mouse model (*32*, *33*); we suggested several mechanisms for the involvement of NO and S-nitrosylation (SNO, for example, NO-mediated posttranslational modification (PTM)) in ASD pathology (*32*–*36*). NO is a small gaseous molecule produced in different organs and tissues, including the central (*37*) and peripheral nervous systems (*38*). At normal physiological levels, NO production and degradation are balanced and involved in normal cell signaling and in regulating numerous physiological functions (*39*, *40*). At higher concentration, however, it becomes toxic and leads to abnormal cell signaling and cell death (*41*). NO can regulate cell signaling by SNO, a form of PTM in which a nitroso group is incorporated into a reactive cysteine thiol to form a nitrosothiol group (*42*). In the nervous system, NO regulates synaptogenesis, vesicle trafficking, neuronal migration, plasticity, and more. Aberrant NO production can cause nitrosative stress by tyrosine nitration of proteins (3-Ntyr) and by forming S-nitrosoglutathione (GSNO), which is a radical recombination between NO and GSH thiyl radicals (RS•); consequently, NO can transnitrosate other thiols on peptides and proteins (*43*). At higher concentrations, NO reacts with superoxide radicals, forms peroxynitrite, and induces cell damage. 3-Ntyr is a product of the reaction of proteins with peroxynitrite. It is an early diagnostic marker of nitrosative stress in neurological disorders (*44*). 3-Ntyr causes disruption of protein native structure and interferes with the phosphorylation ability of tyrosine (*45*). It disrupts many cellular signaling pathways, such as tyrosine nitration of synaptophysin (Syp), which leads to cholinergic dysfunction (*46*, *47*). The free form of 3-Ntyr is equally toxic (*48*).

The dysregulation of NO is responsible for various neurodevelopmental, neuropsychiatric, and neurodegenerative disorders (*33*, *41*). However, the role of NO in ASD remains unknown. To prove the causal effect of NO in ASD pathology, we conducted multidisciplinary experiments using cellular and mouse models as well as clinical samples. Importantly, we found that elevated NO levels contribute to the ASD-phenotype, whereas inhibiting its production via nNOS led to a reversal of it (see Schematic Figure 1). This is the first study to establish a causal link between NO and the development of autism, which may lead to the discovery of novel therapeutic targets for ASD management.

**Fig 1:**
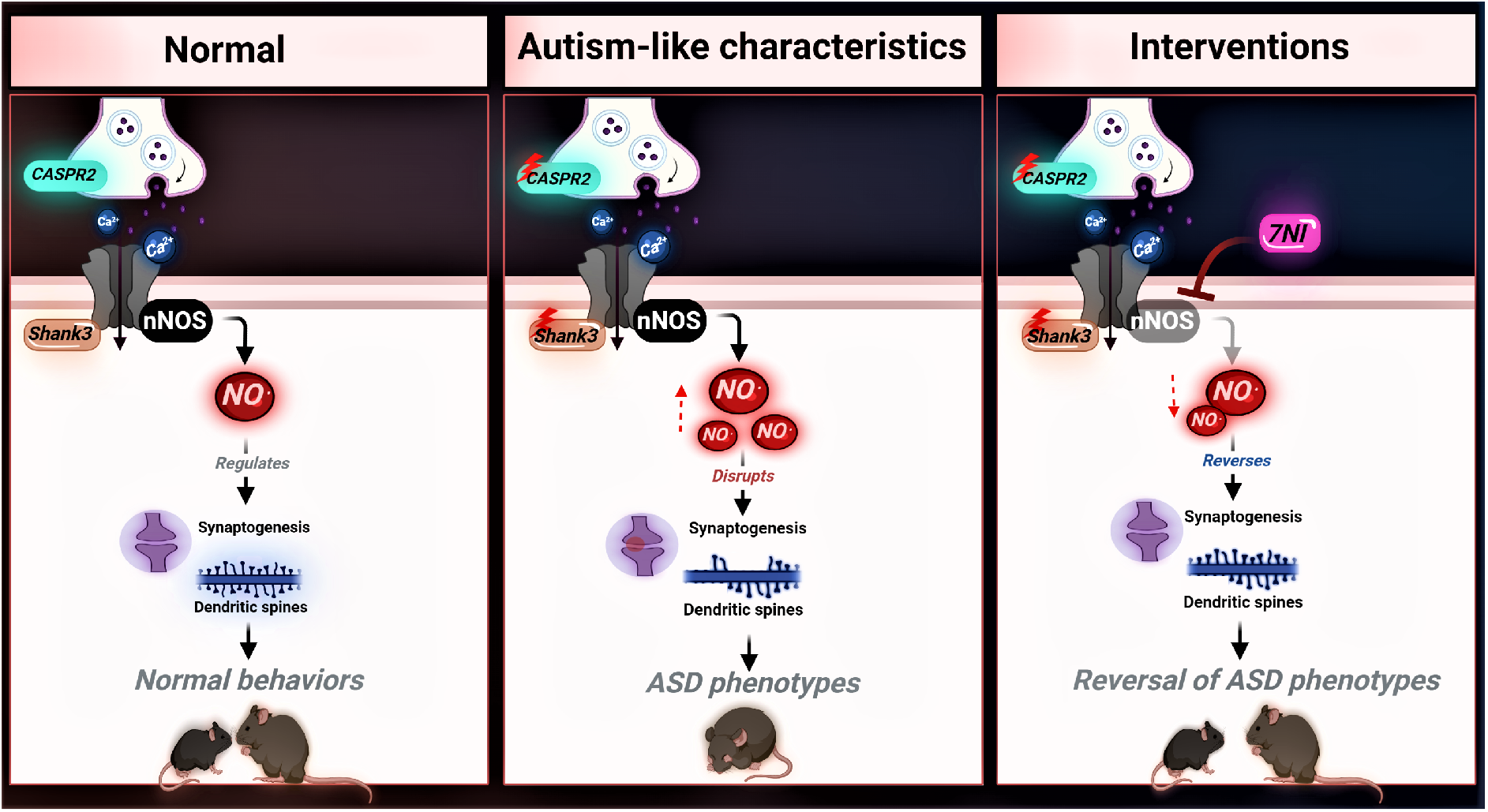
Illustrative figure describing the aberrant role of NO in ASD and the therapeutic potential behind it.

## Results

### NO donor administration to WT mice increased nitrosative stress and reduced synaptogenesis and the expression of glutamatergic and GABAergic proteins

To determine whether NO leads to biochemical and cellular ASD-like phenotypes, we treated juvenile C57BL/6 wild-type (WT) male mice with an NO donor, S-nitroso-N-acetyl penicillamine (SNAP, 20mg/kg) (*49*). SNAP administration increased NO production, which led to 3-Ntyr production in the WT+SNAP group, compared with WT in both the cortex (Fig 2A & 2B) and striatum regions (Fig S1A & S1D). Similarly, the immunofluorescence (IF) analysis of 3-Ntyr in the somatosensory cortex revealed a significant elevation of 3-Ntyr in WT+SNAP mice (Fig 2C & 2G). Next, we determined whether NO affects synaptogenesis. We found that the synaptogenesis biomarker Synaptophysin (Syp) is significantly reduced in SNAP-treated mice, compared with the WT group in both the cortex (Fig 2A & 2B) and in the striatum (Fig S1A & S1D). Furthermore, postsynaptic density protein 95 (PSD95) and Homer levels were decreased in the SNAP-treated group, compared with the WT group in both the cortex (Fig 2A & 2B) and striatum regions (Fig S1A & S1D). Next, we tested the glutamatergic and GABAergic biomarkers that are altered in ASD mouse models. We found a reduction in the NMDA (N-Methyl-D-Aspartate) Receptor 1 (NR1) levels following SNAP treatment in the cortex (Fig 2A & 2B) and striatum (Fig S1A & S1D). IF analysis of NR1 expression in the cortex revealed similar results (Fig 2D & 2G). GABAergic markers such as glutamate decarboxylase 1 (GAD1) and Vesicular GABA transporter (VGAT) were reduced in the SNAP treatment group, compared with WT in the cortex (Fig 2A & 2B) and striatum (Fig S1A & S1D). Furthermore, IF of GAD1 in the somatosensory cortex of mouse brains showed a reduction in the fluorescence intensity in SNAP-treated mice, compared with WT (Fig 2E & 2G). Importantly, we found a significant reduction in the cortical dendritic spine density in SNAP-treated mice (Fig 2F), similarly to the ASD mutant mice as shown below.

**Fig 2.**
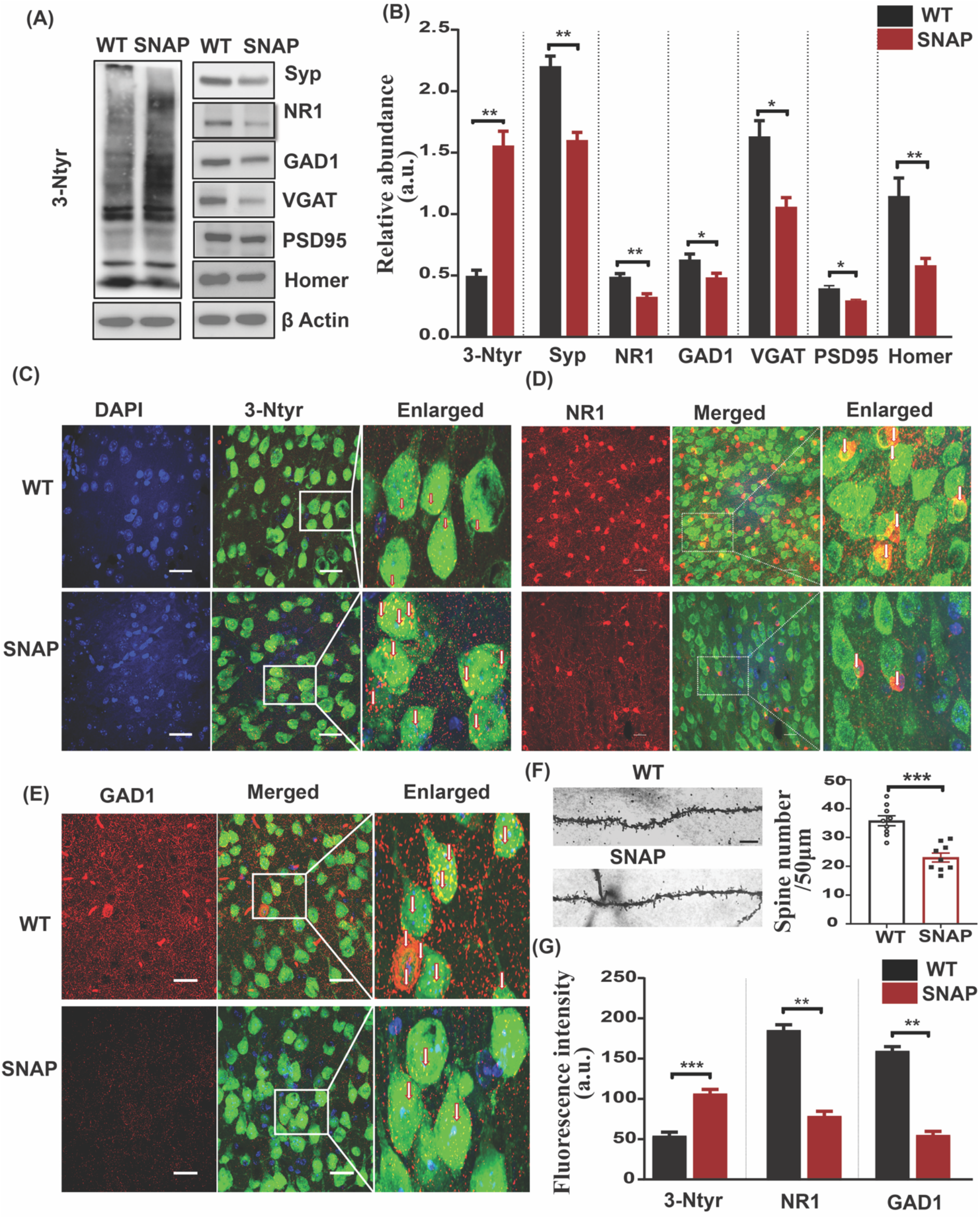
NO donor administration leads to nitrosative stress and synaptic pathology in the cortex of WT mice. **Panel A**: Representative western blots of an indicator of nitrosative stress, 3-Ntyr, and synaptic proteins Syp, NR1, GAD1, VGAT, PSD95, and Homer. β-actin was used as a reference for protein loading. **Panel B:** Statistical analysis of the relative abundance of proteins shown in **Panel A**. **Panel C:** Representative confocal images of 3-Ntyr, NeuN, and DAPI fluorescence. **Panel D:** Representative confocal images of the NR1, NeuN, and DAPI fluorescence. **Panel E:** Representative confocal images of the GAD1, NeuN, and DAPI fluorescence. The enlarged images on **Panels D-F** correspond to the areas shown in the merged images. **Panel F:** Left – representative images of the dendritic spines; right – statistical analysis of the dendritic spine density. **Panel G:** Statistical analysis of the 3-Ntyr, NR1, and GAD1 fluorescence. Student’s two-tailed t-test was used for two-group comparisons in B, F, and G. * p<0.05, ** p<0.01, *** p<0.001. Groups of mice: WT (n=6); SNAP (WT mice treated with the NO donor compound SNAP, n=6).

### NO inhibition in the *Shank3*^Δ*4-22*^ (M1) mouse model led to a reversal in the molecular and synaptic deficits

To determine the involvement of NO in ASD pathology, we inhibited its production pharmacologically in the *Shank3* mouse (M1) model. First, we validated that 3-Ntyr is elevated in M1 mice (Fig 3A & 3C). Administering neuronal nitric oxide synthase (nNOS) inhibitor, 7-nitroindazole (7-NI), (80mg/kg) to the M1 group led to a reversal in the elevated levels of 3-Ntyr in both the cortex (Fig 3A & 3C) as well as the striatum (Fig S1B & S1E). IF of 3-Ntyr in the somatosensory cortex showed similar results (Fig 3D and 3G). In the M1 group, we found a reduction in cortical Syp protein expression, as shown previously (*50*, *51*). 7-NI treatment led to an elevated level of Syp in M1 mice similar to that in the WT mice (which is correlated with synaptogenesis) (*52*) (Fig 3A & 3C). We also found a similar result in the striatum (Fig S1B & S1E). 7-NI administration rescued the downregulation of PSD95 and Homer in the M1 group, compared with WT in the cortical (Fig 3A & 3C) and striatal regions (S1B & S1E). NR1 protein expression was also reduced in the M1 group, compared with WT, whereas 7-NI treatment reversed it in the cortex (Fig 3A & 3C) as well as the striatum (Fig S1B & S1E). IF analysis in the somatosensory cortex revealed the same NR1 protein expression (Fig 3E & 3G). Reduced GAD1 and VGAT levels were normalized by 7-NI administration in the M1 group in the cortex (Fig 3A & 3C) and the striatum (Fig S1B & S1E). IF of GAD 1 in the somatosensory cortex showed similar results (Fig 3F & 3G). Spine density measurement also showed a decrease in the dendritic spine number in the M1 cortical neurons, as found previously (*53*), compared with the WT group, and 7-NI treatment rescued the dendritic spine number in the M1 mice (Fig 3B).

**Fig 3.**
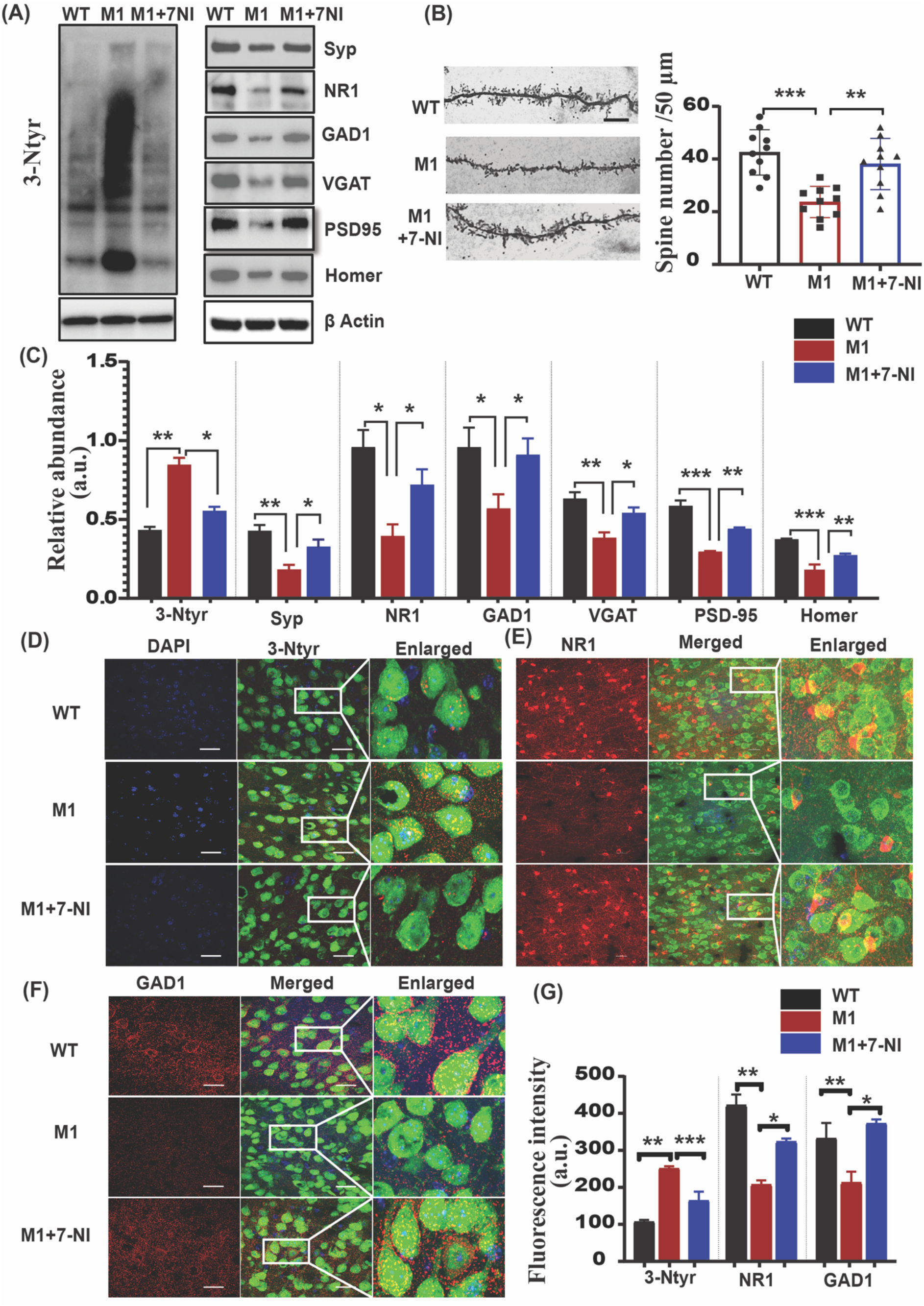
NO inhibition in the *Shank3* mouse model led to a reversal in the molecular and synaptic deficits. **Panel A:** Representative Western blots of an indicator of nitrosative stress, 3-Ntyr; and synaptic proteins Syp, NR1, GAD1, VGAT, PSD95, and Homer. β-actin was used as a reference for protein loading. **Panel B:** Left – representative images of the dendritic spines; right – statistical analysis of the dendritic spine density. **Panel C:** Statistical analysis of the relative abundance of proteins shown in Panel **A**. **Panel D**: Representative confocal images of 3-Ntyr, NeuN, and DAPI fluorescence. **Panel E:** Representative confocal images of the NR1, NeuN, and DAPI fluorescence. **Panel F:** Representative confocal images of the GAD1, NeuN, and DAPI fluorescence. The enlarged images on Panels D-F correspond to the areas shown in the merged images. **Panel G:** Statistical analysis of the 3-Ntyr, NR1, and GAD1 fluorescence. A one-way ANOVA test with the Tukey post hoc test was used for multiple comparisons in **B**, **C,** and **G**. * p<0.05, ** p<0.01, *** p<0.001. Abbreviations, WT (control group), M1 (*Shank3*^Δ*4-22*^), M1+7-NI (*Shank3*^Δ*4-22*^ mice treated with the nNOS inhibitor 7-NI). The number of mice: WT (n=6), M1 (n=6), and M1+7-NI (n=6).

### NO inhibition in the *Cntnap2*^(*-/-*)^ (M2) mouse model led to a reversal in the molecular and synaptic deficits

Our experiments using the SNAP+WT and M1+7-NI groups showed that NO plays a key role in ASD development. To test a larger spectrum and to strengthen our findings, we used another model of ASD, the *Cntnap2*^(*-/-*)^ mutant mouse (M2) model. Similarly to the M1 group, we found that 3-Ntyr protein levels are elevated in the M2 group, compared with WT, and that 7-NI administration reduces its levels in both the cortex (Fig 4A & 4C) and striatum (Fig S1C & S1F) regions. IF analysis of 3-Ntyr in the somatosensory cortex revealed the same findings (Fig 4D & 4G). M2 mice exhibited reduced protein expression of Syp, whereas 7-NI treatment of M2 mice increased Syp protein expression in the cortex (Fig 4A & 4C) and striatum (Fig S1C & S1F). The NR1, GAD1, and VGAT levels were also rescued following NO inhibition by 7-NI in the M2 group in the cortical (Fig 4A & 4C) and striatal regions (Fig S1C & S1F). IF of these proteins in the somatosensory cortex showed similar findings (Fig 4E, 4F, and 4G). Protein expression of Homer and PSD-95 was found to be unchanged in both the cortex (Fig 4A & 4C) and striatum (Fig S1C & S1F). The spine density (the dendritic spine number) was increased following 7-NI treatment in M2 mouse cortical neurons (*54*), compared with the M2 group (Fig 4B).

**Fig 4.**
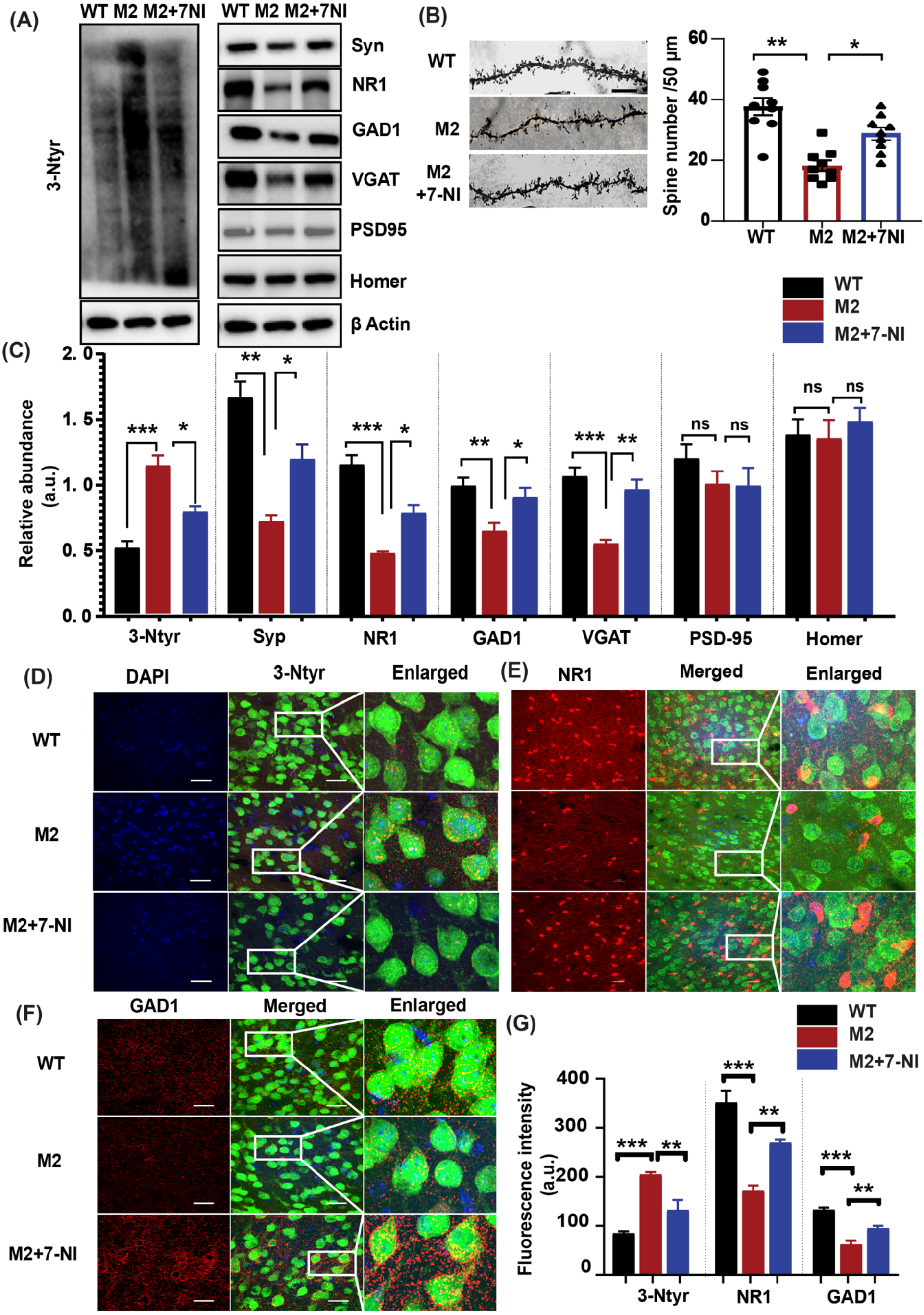
NO inhibition in the *Cntnap2*^(*-/-*)^ mutant (M2) mouse model led to a reversal in the molecular and synaptic deficits. **Panel A:** Representative western blots of an indicator of nitrosative stress, 3-Ntyr, and synaptic proteins Syp, PSD 95, Homer, NR1, GAD1, and VGAT. β-actin was used as a reference for protein loading. **Panel B:** Left – representative images of the dendritic spines; right – statistical analysis of the dendritic spine density. **Panel C:** Statistical analysis of the relative abundance of proteins shown in **Panel A**. **Panel D:** Representative confocal images of 3-Ntyr, NeuN, and DAPI fluorescence. **Panel E:** Representative confocal images of the NR1, NeuN, and DAPI fluorescence. **Panel F:** Representative confocal images of the GAD1, NeuN, and DAPI fluorescence. The enlarged images on Panels **D-F** correspond to the areas shown in the merged images. **Panel G:** Statistical analysis of the 3-Ntyr, NR1, and GAD1 fluorescence. A one-way ANOVA test with the Tukey post hoc test was used for multiple comparisons in **B**, **C,** and **G**. * p<0.05, ** p<0.01, *** p<0.001. Abbreviations, WT (control group), M2 (*Cntnap2*^(*-/-*)^), M2+7-NI (*Cntnap2*^(*-/-*)^ mice treated with the nNOS inhibitor 7-NI). The number of mice: WT (n=6), M2 (n=6), and M2+7-NI (M2 mice treated with the nNOS inhibitor 7-NI; n=6).

### NO inhibition reversed the ASD-like behavior in the *Shank3*^Δ*4-22*^ mouse model

To correlate our biochemical and cellular findings with the behavioral side, we tested the potential effects of inhibiting NO on the ASD-like behavior in M1 mice. In brief, M1 mice were treated with 7-NI (80mg/kg) intraperitoneal (IP), and WT mice were treated with their respective vehicle. Initially, we performed the open field test to examine the motor activity in mice (the distance traveled and the velocity). In the motor activity test, we did not see any changes between the three groups (Fig 5A). To test anxiety-like behavior, we conducted an elevated plus maze test. The analysis revealed that the M1 mice spent significantly more time in closed arms than the WT mice, indicating increased anxiety, whereas the M1+7-NI mice spent less time in closed arms, and exhibited less anxiety-like behavior (Fig 5B).

**Fig 5.**
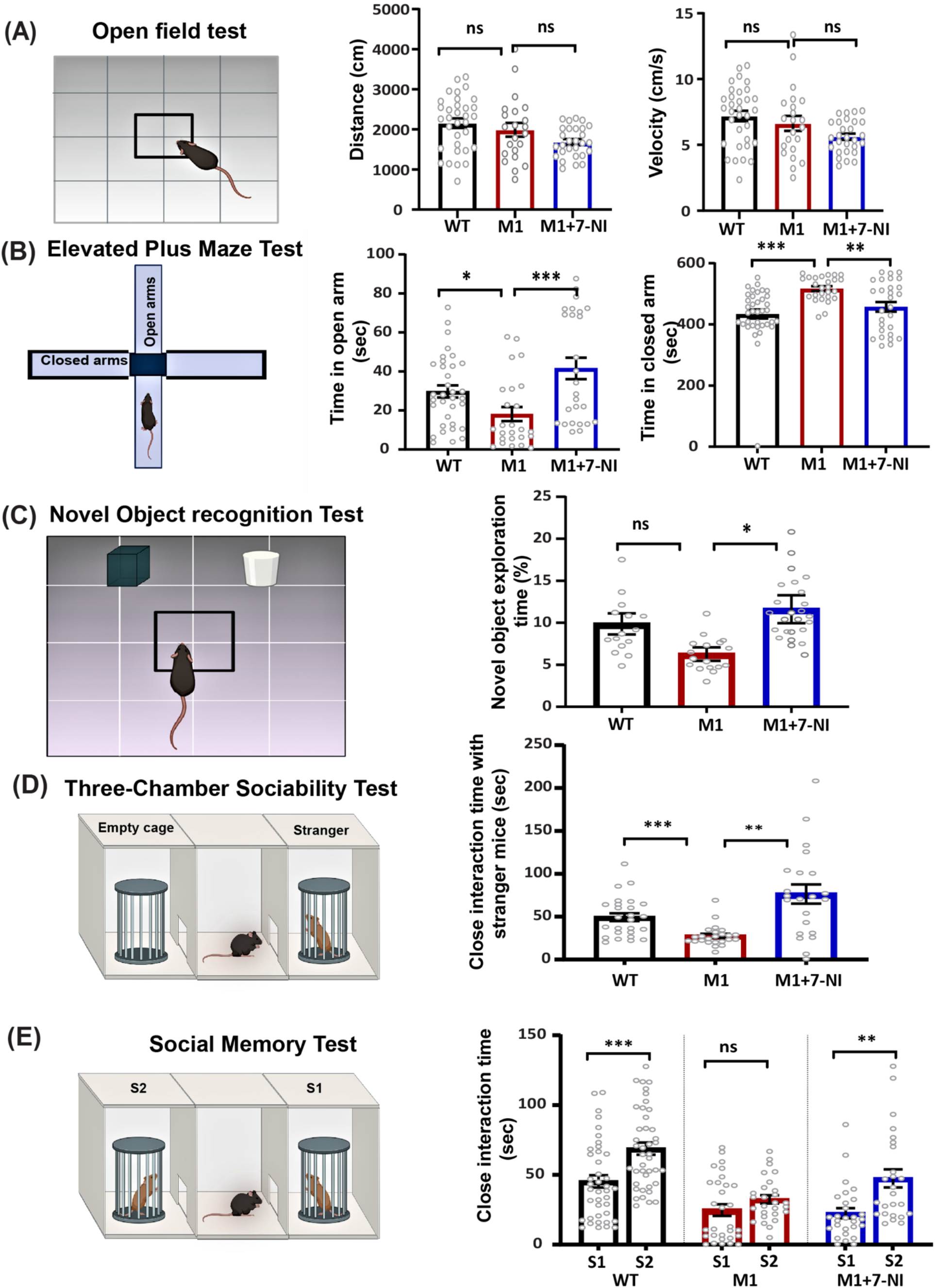
7-NI-treatment reversed autistic behavior abnormalities *Shank3^-/-^* mutant mice. **Panel A:** Left-An illustrative representation of the motor activity test, Right-Statistical analysis of the distance traveled and the velocity in the WT (n=35), M1 (n=22), and M1+7-NI groups (n=28). **Panel B:** Left-An illustrative representation of the elevated plus maze test, Right-Statistical analysis of the total time spent in open arms and closed arms by the WT (n=37), M1 (n=26), and M1+7-NI group (n=28). **Panel C:** Left-An illustrative representation of the NOR test, Right-Statistical analysis of the time spent interacting with a novel object by the WT (n=17), M1 (n=19), and M1+7-NI (n=25) groups. **Panel D:** Left-An illustrative representation of the three-chamber sociability test, Right-Statistical analysis of the close interaction time with stranger mice by the WT (n=32), M1 (n=24), and M1+7-NI group (n=22). **Panel E:** Left-An illustrative representation of the social memory test, Right-Statistical analysis of the close interaction time with a novel mouse by the WT (n-52), M1 (n=30), and M1+7-NI group (n=28). A one-way ANOVA test with the Tukey post hoc test was used for multiple comparisons in all tests. * p<0.05, ** p<0.01, *** p<0.001, ns=not significant. Abbreviations, WT (control group), M1 (*Shank3*^Δ*4-22*^), M1+7-NI (*Shank3*^Δ*4-22*^ mice treated with the nNOS inhibitor 7-NI). S1=Familiar mouse, S2= novel mouse.

To test novelty-seeking behavior, we conducted a novel object recognition (NOR) test. The results indicated that the M1 mice treated with 7-NI spent more time exploring the novel object, compared with the M1 group, reflecting a reversal in the deficits of novelty-seeking behavior. (Fig 5C). The three-chamber sociability test showed that the M1 mice spent significantly less time interacting with the stranger mouse, compared with their WT littermates, indicating deficits in social behavior, whereas the M1+7-NI mice spent more time interacting with the stranger mouse, compared with their untreated counterparts M1 (Fig. 5D). Next, we performed a social memory test. In the social memory test, the M1 mice showed no interest in interacting with the novel mouse, whereas the M1+7-NI mice spent more time interacting with the novel mouse, compared with the familiar mice (Fig 5E). The behavioral results suggest that the autistic behavior phenotypes observed in the M1 mice can be reversed by inhibiting neuronal NO production.

### NO inhibition reversed the ASD-like behavior in the *Cntnap2*^(*-/-*)^ (M2) mouse model

Following our behavioral findings using the M1 mouse model, we tested the same behavioral parameters in the M2 mouse group. The open field test revealed a non-significant difference in the distance traveled and the velocity between the three tested groups (Fig S2A). The NOR test showed that the M2 mice spent significantly less time exploring the novel object than the WT mice, reflecting reduced novelty seeking and restricted interests. However, following treatment with 7-NI (M2+7-NI), the mice exhibited a significant preference for the novel object over the familiar one (Fig 6A). In the elevated plus maze, the analysis revealed that the M2 mice spent significantly more time in closed arms than the WT mice, indicating increased anxiety, whereas the M2+7-NI mice spent less time in closed arms (Fig 6B). Next, we performed a three-chamber sociability test that showed that the M2 mice spent significantly less time interacting with the stranger mouse than their WT littermates, indicating deficits in social behavior, whereas the M2+7-NI mice spent a longer time interacting with the stranger mouse, compared with their untreated counterparts M2 (Fig 6C). The social memory test demonstrated that the WT mice spent significantly more time interacting with the novel mouse (S2), compared with the familiar one (S1); in contrast, no significant difference was observed among the M2 and the M2+7-NI mice (Fig S2B). All together, the results suggest that a mutation in the *Cntnap2* gene leads to autistic phenotypes, inhibits NO production, and attenuates ASD behavioral phenotypes.

**Fig 6.**
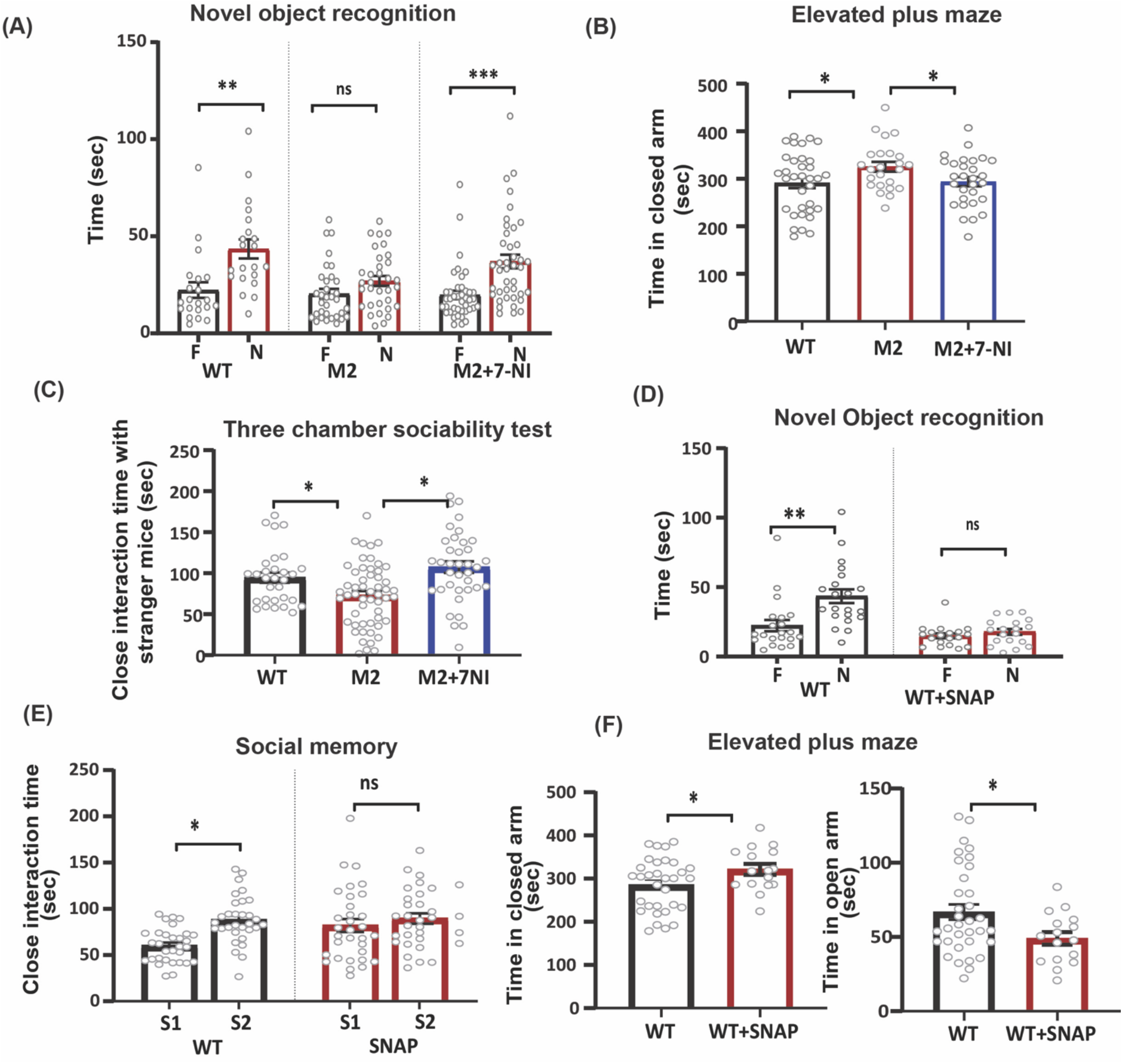
7-NI-treatment corrects autistic behavior abnormalities in *Cntnap2*^(*-/-*)^ mutant mice and SNAP treatment increases behavioral abnormalities in WT mice. **Panel A:** Statistical analysis of the time spent interacting with a novel object by the WT (n=33), M2 (n=41), and M2+7-NI (n=21) groups. **Panel B**: Statistical analysis of the total time spent in open arms and closed arms by the WT (n=34), M2 (n=24), and M2+7-NI (n=29) groups. **Panel C:** Statistical analysis of close interaction time with stranger mice by the WT (n=54), M2 (n=38), and M2+7-NI (n=34) groups. **Panel D:** Statistical analysis of time spent interacting with a novel object by the WT (n=21) and SNAP treatment groups (n=19). **Panel E:** Statistical analysis of close interaction time with the novel mouse by the WT (n=33) and SNAP treatment groups (n=32). **Panel F:** Statistical analysis of elevated plus maze in the WT (n=34) and SNAP treatment groups (n=16). A one-way ANOVA test with the Tukey post hoc test was used for multiple comparisons in A, B, and C. Student’s twotailed t-test was used for two-group comparisons in D, E, and F. * p<0.05, ** p<0.01, *** p<0.001, ns= not significant. Abbreviations, WT (control group), M2 (*Cntnap2*^(*-/-*)^), M2+7-NI (*Cntnap2*^(*-/-*)^ mice treated with the nNOS inhibitor 7-NI). SNAP (WT mice treated with the NO donor compound SNAP). S1=Familiar mouse, S2=novel mouse. F= Familiar object, N=Novel object.

### NO donor administration induced ASD-like behavior in WT mice

A large set of behavioral tests were conducted following the treatment of NO donor, SNAP (20mg/kg), to WT mice, to examine the potential effects of high levels of NO on their behavior. The motor activity test analysis revealed no significant difference in the distance traveled and the velocity between the WT and the SNAP-treated mice (Fig S2D). Furthermore, we conducted the NOR test in the WT and SNAP-treated mice, which showed that the WT mice spent significantly more time exploring the novel object than the familiar object. In contrast, the SNAP-treated mice failed to display a preference for the novel object, indicating a lack of interest and reduced novelty seeking (Fig 6D). Furthermore, a three-chamber sociability test was conducted. In the first session, we did not find a significant difference between the WT and SNAP-treated mice regarding the time spent with stranger mice or while in an empty cage. However, the second session (social memory) showed that the SNAP-treated mice spent less time interacting with the novel mouse than with the familiar one (S1), indicating reduced social memory (Fig 6E). Next, in the elevated plusmaze test, the analysis revealed that the SNAP-treated mice spent more time in closed arms, compared with WT, reflecting an anxiety-like behavior (Figure 6F). All together, the data suggest that a high level of NO induces ASD-like behavior.

### NO inhibition promoted synaptogenesis and reduced nitrosative stress in primary cortical neurons derived from mutant ASD mice

To validate our *in-vivo* experiments at the cellular level and confirm that neurons are specifically involved in the NO-mediated pathological mechanism, we conducted a set of experiments using primary cortical neuronal cell cultures. We selectively cultured cortical neurons from the embryos of WT and both mutant groups and treated them with vehicle and 7-NI (20uM), respectively (see the detailed MTT assay in Supplementary Table 1). Primary neurons from both mutant groups showed elevated 3-Ntyr levels, whereas treatment with 7-NI reduced the expression of 3-Ntyr in both mutant groups (Fig 7A-B, 7E-F), which is consistent with the in-vivo findings. To examine synaptogenesis, we further investigated the expression of Syp in both mutant groups. Syp expression is significantly downregulated in the mutant group, whereas 7-NI treatment rescued its expression towards normalcy (Fig 7C-D, 7G-H).

**Fig 7.**
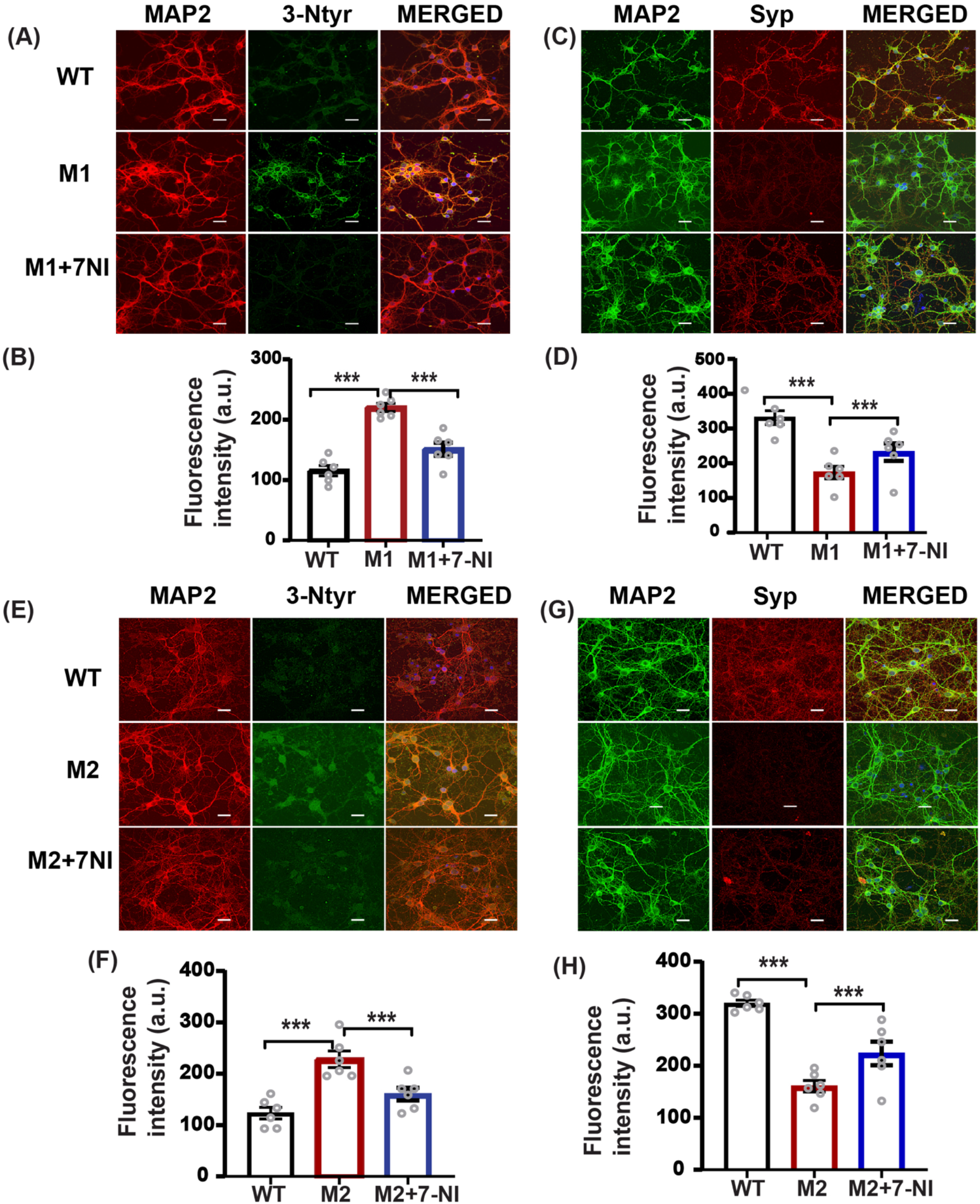
NO contributes to nitrosative stress and reduced synaptogenesis in the primary cortical neurons of the *Shank3^-/-^* and *CNTNAP*^*-/-*^ mutant groups. **Panel A:** Representative confocal images of the MAP2, 3-Ntyr, and DAPI in WT, M1, and M1+7-NI. **Panel B:** Statistical analysis of the Ntyr fluorescence. **Panel C:** Representative confocal images of the Syp, MAP2, and DAPI fluorescence in WT, M1, and M1+7-NI. **Panel D:** Statistical analysis of the Syp fluorescence. **Panel E:** Representative confocal images of the MAP2, 3-Ntyr, and DAPI in WT, M2, and M2+7-NI. **Panel F:** Statistical analysis of the Ntyr fluorescence. **Panel G:** Representative confocal images of the Syp, MAP2, and DAPI fluorescence in WT, M2, and M2+7-NI. **Panel H:** Statistical analysis of the Syp fluorescence. A one-way ANOVA test with the Tukey post hoc test was used for multiple comparisons in all groups. * p<0.05, ** p<0.01, *** p<0.001. Abbreviations, WT (primary neurons isolated from WT embryos), M1 (primary neurons isolated from *Shank3*^Δ*4-22*^ embryos), M1+7-NI (primary neurons isolated from *Shank3*^Δ*4-22*^ embryos treated with the nNOS inhibitor 7-NI). M2 (primary neurons isolated from *Cntnap2*^(*-/-*)^ embryos), M2+7-NI (primary neurons isolated from *Cntnap2*^(*-/-*)^ embryos treated with the nNOS inhibitor 7-NI). WT (n=6), M1 (n=6), and M1+7-NI (n=6), M2 (n=6), M2+7-NI (n=6).

### nNOS affects 3-Ntyr and Syp in human cell lines

To translate our animal research into human results, we conducted a set of experiments using human neuroblastoma cells and human plasma samples. Our findings on primary neuronal culture and a selective nNOS inhibition study suggest that nNOS is the main isoform among the three NOS isoforms involved in the pathology. However, in order to validate this, we knocked down the expression of *SHANK3* and *nNOS* in differentiated neuroblastoma cells. First, we tested 3-Ntyr expression and found that *SHANK3* knockdown increased the expression of 3-Ntyr, whereas double knockdown of *SHANK3* and nNOS reduced the expression of 3-Ntyr proteins (Fig 8A and 8B). In the experiments that followed, we evaluated the expression of Syp and found that *SHANK3* knockdown reduced its expression, whereas double knockdown with *nNOS* and *SHANK3* rescued its expression towards normalcy (Fig 8C & 8D).

**Fig 8.**
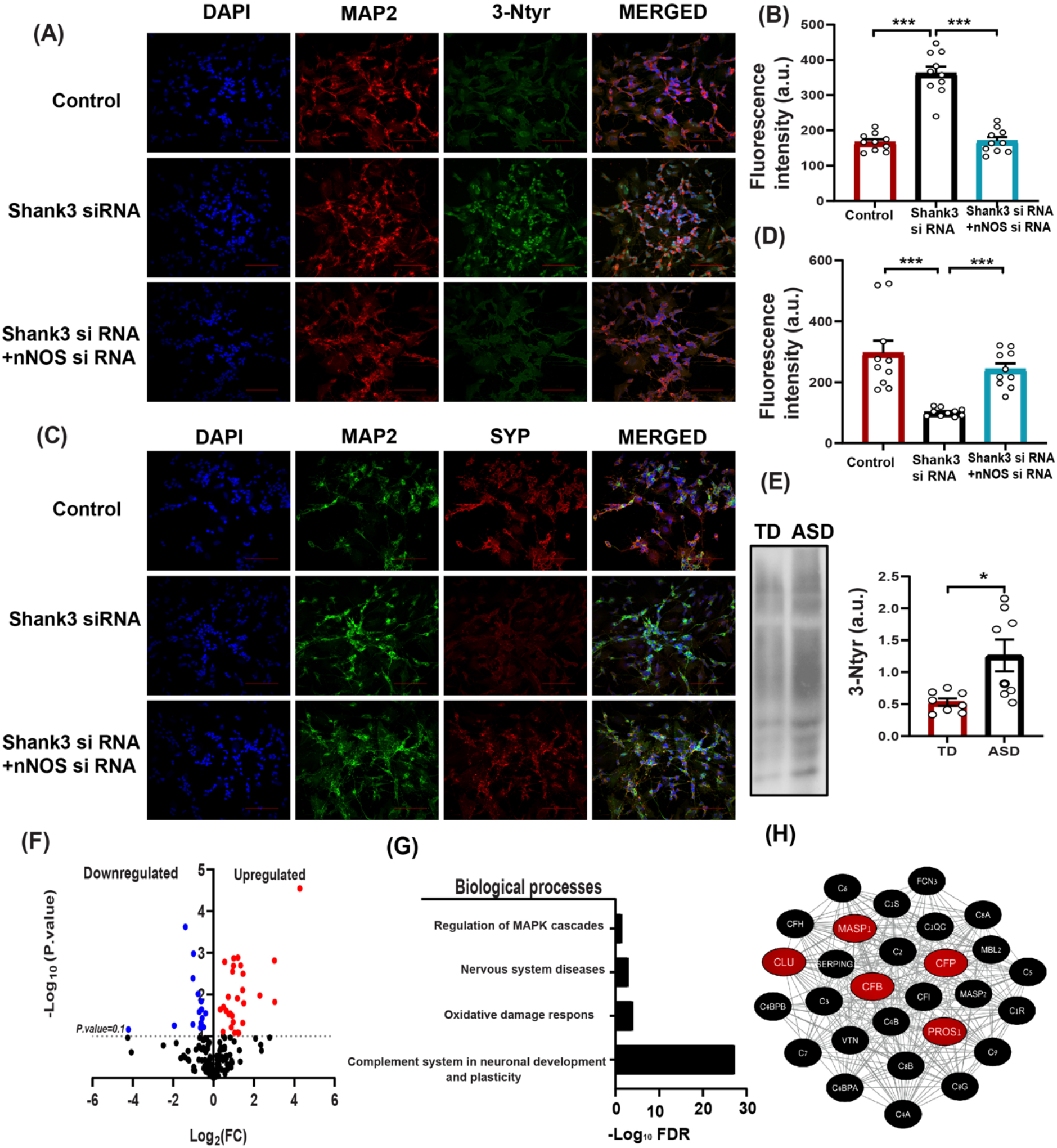
NO reprogramming in human cell lines and blood samples from ASD children. **Panel A:** Representative confocal images of MAP2, 3-Ntyr, and DAPI in SHSY5Y (n=10); *SHSY5Y*+*siSHANK3* (n=10); and *SHSY5Y*+*siSHANK3*+*si-nNOS* (n=10). **Panel B:** Statistical analysis of the 3Ntyr fluorescence. **Panel C**: Representative confocal images of the Syp, MAP2, and DAPI fluorescence in SHSY5Y (n=10); *SHSY5Y*+*siSHANK3* (n=10); and *SHSY5Y*+*siSHANK3*+*si-nNOS* (n=10)). **Panel D**: Statistical analysis of the Syp fluorescence. **Panel E**: Representative western blot of 3-Ntyr in human plasma samples in TD (n=8), and ASD (n=8) with statistical analysis. **Panel F**: Volcano plot displaying the log _2_ (FC) on the x-axis plotted against the −log _10_ (p-value) on the y-axis for all the identified proteins that were differentially expressed in the plasma of TD (n=20) and ASD (n=19) individuals. Proteins that were significantly upregulated in ASD (P <0.1) are denoted in red, whereas the proteins that were significantly downregulated are denoted in blue (p<0.1). **Panel G**: BP analysis was conducted on the identified plasma SNO proteins using STRING, version 11.5. Each bar represents the −log _10_ (FDR). **Panel H**: Clustering analysis of the plasma SNO-proteins. “The complement system in neuronal development and plasticity” was enriched (the number of proteins=28, FDR=6.26e-28). The red nodes are the proteins that were significantly changed between the ASD and TD individuals (p<0.1). STRING, version 11.5, and Cytoscape software version 3.3.0 was used to generate this figure. Student’s twotailed t-test was used for two-group comparisons in **E**. A one-way ANOVA test with the Tukey post hoc test was used for multiple comparisons in **B** and **D**. * p<0.05, ** p<0.01, *** p<0.001. Abbreviations, ASD=low functioning ASD patients, TD=typically developed volunteers, FDR=false discovery rate

### Elevation of nitrosative stress biomarker and reprogramming the SNO-proteome in the blood samples of ASD children

Finally, we measured the levels of 3-Ntyr and analyzed the SNO-proteome in human blood samples. We tested 3-Ntyr expression in blood plasma taken from 20 typically developed (TD) individuals and 19 autistic individuals (ASD) (ages: 2-8 years) (see supplementary Table 2 for detailed clinical data). We randomly picked 8 ASD and 8 TD blood samples to conduct the WB analysis. We found an increased expression of 3-Ntyr in ASD, compared with the TD children (Fig 8E). To identify the SNOed proteins, we used the novel SNOTRAP-based mass-spectrometry technology developed recently (*32*, *55*). We mapped the SNO-proteome in both groups and identified 171 proteins that were differentially SNOed in the blood of autistic individuals (for the IDs of SNO-proteins in the plasma samples, see supplementary Table 3). Volcano plot analysis was conducted to quantify and visualize the changes in the expression of the SNO proteins in the blood samples of both groups. Importantly, the analysis revealed a total of 28 proteins that were significantly upregulated (red dots) in the blood of ASD individuals and a total of 15 proteins that were significantly downregulated (blue dots) in the blood of ASD individuals (Fig 8 F).

To obtain a system-level insight into the SNO-protein functionalities, we applied a system biology analysis by dissecting the biological processes (BP) and the pathways that might be affected due to the altered pattern of the SNO (for details on the system biology analysis, see Supplementary Table 4). BP analysis revealed the enrichment of brain-related processes and pathways, which includes nervous system diseases (FDR= 0.0012), regulation of MAPK cascades (FDR= 0.0388), complement systems in neuronal development and plasticity (FDR= 6.26e-28), as well as oxidative damage response (FDR= 0.00012) (Fig 8G).

We applied functional clustering analysis to get further functional insights into the data. We used the gene ontology (GO) classification system to classify the proteins into different clusters based on their enriched biological processes. Twenty-eight of the identified SNO-proteins formed a distinct functional cluster related to the complement system in neuronal development (FDR= 6.26e-28) and plasticity, which is correlated with the BP analysis. Further analysis should be done to validate the link between NO and complement systems in ASD pathology (Fig 8H).

## Discussion

Our study is designed to examine the effect of NO on the development of ASD. We found strong evidence of its involvement in ASD pathology. Importantly, we confirmed this in cellular and animal models as well as clinical samples. Since the molecular mechanisms underlying ASD pathogenesis remain largely unknown, we provided data that show the NO role in ASD pathology at the molecular, cellular, and behavioral levels. An increase of Ca^2+^ influx in ASD pathology, including in human and mouse models of *Shank3* and *Cntnap2*, has already been reported (*25*, *56*, *57*). Ca^2+^ activates nNOS, which further leads to massive production of NO (*58*). Aberrant NO production induces oxidative and nitrosative stress, leading to increased 3-Ntyr production and aberrant protein SNO. Our data showed an increase in 3-Ntyr production in both mouse models (*Shank3*^Δ*4-22*^, *Cntnap2*^(*-/-*)^ and in human blood plasma. The elevated levels of 3-Ntyr in our study is consistent with previous postmortem examinations of ASD patients showing the accumulation of this molecule in the brain (*59*).

Abnormal synaptogenesis is a key factor in ASD pathology. One of the biomarkers of synaptogenesis is Syp (*60*, *61*), which is prone to nitrosative stress, and NO regulates its expression (*62*). It was found that when Syp is nitrated, this leads to proteasomal degradation (*63*, *64*). In our study, we found that administering NO donor reduces the expression of Syp as well as the high levels of NO in *Shank3*^Δ*4-22*^ and *Cntnap2*^(*-/-*)^ mutant mice, leading to similar results. Administering 7-NI in both mutant models restored the expression of Syp, which directly confirms that NO plays a key role in synaptogenesis in ASD. Dendritic spines are tiny membranous outgrowths that are the primary sites for excitatory synaptic transmission (*65*). Spine density measurement also showed a reduction of dendritic spines in NO donor-treated mouse cortical neurons, compared with the WT group, whereas 7-NI treatment in mutant mice restored its number. In the CNS, dendritic spines affect synaptic connectivity, which is essential for neuronal circuits (104, 105) and learning (*66*). Dysregulation of dendritic spine formation has been reported in ASD patients (*67*, *68*). Therefore, the restoration of dendritic spine density following 7-NI treatment indicates that NO is a critical factor in the development of the synapthopathology of ASD.

Nitrosative stress also disrupts GABAergic and glutamatergic signaling in the brain (*69*, *70*); S-nitrosylation of NR1 (*71*) and GAD65 (*72*) leads to their degradation. It is in agreement with our results that NO donor leads to the downregulation of the GABAergic and glutamatergic markers. The mutant ASD mice with elevated NO levels in the brain also showed similar results. Administering 7-NI in both mutant models restored the glutamatergic and GABAergic markers, which indicated that NO plays a role in disrupting the glutamatergic and GABAergic systems in ASD.

Behavioral studies revealed that nitrosative stress induces behavioral abnormalities in rodents (*73*, *74*). Similar to both the *Shank3* (*17*, *75*, *76*) and *Cntnap2* (*25*, *77*, *78*) mouse models, WT mice treated with NO donor exhibited a reduced interest in novel objects, as well as impaired sociability and increased anxiety. However, administering 7-NI led to a reversal in most of the behavioral phenotypes in mutant mice. These results suggest that high NO concentrations in ASD pathology could potentially induce behavioral and cognitive deficits. Collectively, we concluded that inhibition of NO production leads to the reversal of ASD-like behavior.

In addition, we selectively cultured primary cortical neurons from both mutant groups to rule out the effects of (iNOS) or endothelial NOS (eNOS) in NO production due to *SHANK3* and *CNTNAP2* mutations. Thus, similar to the *in-vivo* results, we found a loss of Syp expression and elevated levels of 3-Ntyr. Treating these cells with selective nNOS inhibitor,7-NI, significantly restored these parameters. This result confirms the involvement of nNOS in aberrant NO production leading to molecular and cellular deficits. Similarly, genetic manipulation of nNOS in *SHANK3* knocked-down human cell lines showed a reversal in 3-Ntyr and synaptic defects. Collectively, our targeted pharmacological and genetic manipulations of nNOS showed its involvement in ASD.

Bioinformatics analysis of the SNO-proteome of human blood samples revealed an enrichment of key BPs that are associated with ASD progressions, such as regulation of MAPK cascades (*79*), nervous system diseases (*80*), a complement system in neuronal development and plasticity (*81*), and oxidative damage response (*82*), which is consistent with the above references. GO classification analysis of the SNO-proteins showed the involvement of complement system proteins (CFP, CFB, CLU, PROS1, and MASP1) in ASD plasma samples. Previous studies have demonstrated the involvement of complement systems in ASD (*83*–*85*). NO also plays a role in complement system activation (*86*, *87*). However, to study the role of NO in complement systems in ASD, further investigation is needed.

Collectively, the results of our study show for the first time that NO plays a key role in ASD development. We found that NO can affect the synaptogenesis, glutamatergic, and GABAergic systems in the cortex and striatum, which converge into ASD-like behavioral deficits. This work implies that NO is an important pathological factor in ASD and that it will open novel future directions to examine NO in diverse mutations on the spectrum as well as other neurodevelopmental disorders and psychiatric diseases. Finally, this is the first study to establish a causal link between NO and ASD, leading to the discovery of novel NO-related drug targets for the disorder and suggesting nNOS as a precise target for treatment.

## Material, methods, and participants

### Materials

Primary antibodies, Anti-GAD1 (#41318), Anti-NeuN (#94403), Anti-β-Actin (#3700), Anti-PSD95 (#36233), Anti-Caspr2 (#3731), Anti-nNOS (#4231), Anti-synaptophysin (#36406) and secondary antibodies, Anti-Rabbit Alexa fluor 594 (#8889), Anti-mouse (Alexa Fluor 488 (#4408), HRP conjugated anti-rabbit (7076S), HRP conjugated anti-mouse (7074S) and ProLong gold Antifade with DAPI (#8961), protease phosphatase inhibitor cocktail (#5872) were purchased from Cell Signaling Technology (Danvers, MA, USA). Anti-3Ntyr antibody (ab110282), Anti-homer 1 (ab184955), Anti-NMDAR1 antibody (ab109182), Anti-SLC32A1/ VGAT antibody (1b235952), and anti-MAP2 (ab11268) were purchased from Abcam, Cambridge, UK. Anti-Shank 3 (sc-377088) and Anti-NOS 2 (sc7271) were purchased from Santa Cruz Biotechnology, Dallas, TX, USA. Other general chemicals were purchased from Sigma Aldrich (St. Louis, MO, USA) and Bio-Rad Laboratories (Haifa, Israel).

### Animal housing, pharmacological interventions, and tissue dissection

All animal experiments were conducted under the guidelines of the Institutional Animal Care and Use Committee and the Association for Assessment and Accreditation of Laboratory Animal Care International. Juvenile (6–8-week-old) male *Shank3*^(*-/-*)^ (4-22 exon deletion) (*88*) and their control littermates (used as WT/control for the *Shank3* mice), as well as *Cntnap2*^(*-/-*)^, (*25*) and Wild-Type (WT) C57BL/6 mice were used in this study. The C57BL/6 mice served as WT mice for the *Cntnap2* mice. All mice were purchased from the Jackson Laboratory (Farmington, CT, USA). The animals were kept at a room temperature (RT) of 23°C in a 12-hour light/dark cycle and fed ad libitum with standard mouse chow and water.

The pharmacological interventions were performed by daily intraperitoneal (IP) injection for 10 consecutive days. An NO donor, SNAP, was administered to the WT mice at 20 mg/kg (*49*). A selective nNOS inhibitor, 7-NI, was injected into *Shank3*^Δ*4-22*^ mutant and *Cntnap2*^(*-/-*)^ mice at a dose of 80 mg/kg (*89*). Thus, the following groups of mice were used in this study: (1) WT+vehicle, (2) WT+SNAP, (3) *Shank3*^Δ*4-22*^(M1), (4) *Cntnap2*^(*-/-*)^ (M2), (5) *Shank3*^Δ*4-22*^+7-NI (M1+7-NI), and (6) *Cntnap2*^(*-/-*)^+7-NI (M2+7-NI). Mice were randomly selected either for the behavioral or biochemical tests. After the 10th injection of treatment or vehicle, the animals were euthanized. For western blot analysis, the entire cortex and striatum were isolated, snap-frozen in liquid nitrogen, and stored at −80°C for further use, as described previously (*34*). For IF, mice were intracardially perfused with 0.9% saline and 4% paraformaldehyde. The whole brain was isolated and stored in 10% paraformaldehyde for 24 h. Thereafter, the brains were gradually dehydrated with 10%, 20%, and 30% sucrose gradients. Cryostat (Leica company, Wetzlar, Germany) was used to cut the brain sections.

### Western blots

Tissues were homogenized in freshly prepared RIPA buffer (30mM HEPES, pH 7.4, 150mM NaCl, 1% Nonidet P-40, 0.5% sodium deoxycholate, 0.1% sodium dodecyl sulfate, 5mM EDTA, 1mM NaV04, 50mM NaF, 1mM PMSF, 1% protease, and phosphatase inhibitors cocktail, pH 7.7) on ice using Teflon pestle and a Jumbo Stirrer (Thermo Fisher Scientific, Waltham, MA, USA). The homogenates were centrifuged at 13000xg for 30 min at 4°C. The supernatant was collected and the protein concentration was estimated by Bicinchoninic Acid (BCA) Protein Assay (Sigma Aldrich). The samples were subjected to polyacrylamide gel electrophoresis, followed by wet transfer onto a polyvinylidene fluoride (PVDF) membrane (Bio-Rad Laboratories). Non-specific sites were blocked by either 5% dried skimmed milk or 5% bovine serum albumin (BSA) in Tris-buffered saline with Tween 20 (TBST), containing 135 mM NaCl, 50 mM Tris, and 0.1% Tween 20, pH 7.4, for 2 hours at room temperature. PVDF membranes containing transferred proteins were incubated with a primary antibody overnight at 4°C under shaking conditions. The following primary antibodies (were used: anti-Syp (1:1000), anti-Homer (1:1000), anti-NR1 (1:1000), anti-GAD1 (1:1000), anti-nNOS (1:1000), anti-PSD95 (1:1000), anti-iNOS (1:1000), anti-VGAT (1:1000), anti-β-Actin (1:1000), anti-3-Ntyr (1:1000), and anti-Shank 3 (1:1000). After exposure to primary antibodies, the membranes were washed with TBST and incubated with anti-mouse/rabbit HRP secondary antibody for 1 h at room temperature. Specific binding of the protein of interest was detected using ECL substrate (Bio-Rad Laboratories). The bands were visualized using the Bio-Rad Chemidoc imaging system (Hercules, CA, USA).

### Cell culture

Human neuroblastoma (SH-SY5Y) cells were obtained from the American Type Culture Collection (Manassas, VA, USA) and maintained in a 1:1 mixture of Ham’s F-12 and Dulbecco’s modified Eagle’s medium (DMEM) supplemented with 10% fetal bovine serum, 2 mM L-glutamine, and 1% penicillin-streptomycin in a humidified atmosphere at 37°C and 5% CO_2_. Cells were plated onto a non-coated 35 mm dish at a density of 6.0 × 10^4^ cells/cm^2^ for western blotting and immunocytochemistry.

### SH-SY5Y Differentiation

SH-SY5Y cell differentiation was carried out by applying Retinoic acid (RA) and BDNF simultaneously, as described previously (*90*). In brief, the cells in passage numbers 6-8 were used in this experiment and kept under differentiation media over a period of 11-12 days. The media were changed every alternate day. Then, the differentiated cells were further used in the experiment (*91*).

### siRNA transfection

SH-SY5Y cells were cultured at a density of 6.0 × 10^4^ cells/cm^2^ and transfected with 20nM of SHANK3 and 25nM of nNOS siRNA (Life Technologies) simultaneously for 24 h, using Lipofectamine RNAi MAX (#13778030, Life Technologies) according to the manufacturer’s protocol. Then, the media was changed after 6 h, and cells were allowed to grow for 72h. Finally, the cells were washed with PBS and harvested for western blotting, and cells cultured on a coverslip were used for confocal microscopy (*90*, *92*).

### Primary cortical neuronal cultures

Primary cortical neurons were isolated from fetal mouse brains on embryonic days 16-17. In brief, the cortex isolated from embryonic mouse brains was placed in DMEM (Thermo Fisher Scientific) and incubated with 0.25% trypsin + 0.02% EDTA (Sigma-Aldrich) at 37°C for 15 min. Neurons were mechanically dissociated by pipetting and seeded on a poly-D-lysine-coated glass (Sigma-Aldrich) in 6 well plates. The density was 3 × 10^5^ in 6 wells on 22mm PDL-coated slides for microscopic observation. Cells were cultured in neurobasal medium (Gibco) containing 2% B27 supplement, and 1% penicillin-streptomycin, in a humidified incubator at 37°C, 95% air, and 5% CO_2_. On the second day of neuronal culture, half the media volume was replaced with the same volume of fresh neurobasal media with 2uM cytosine-β-D-arbinofuranoside (Sigma-Aldrich) to avoid glial cell proliferation. The half media was changed with new media every 3^rd^ day. Cytosine-β-D-arbinofuranoside was treated only until the 7^th^ day, and the cells were grown for 14 days (*93*).

### MTT cell viability assay

The effects of 7-NI on primary cortical neurons were determined using a colorimetric 3-(4,5-dimethylthiazol-2-yl)-2,5-diphenyl-tetrazolium (MTT) assay (Sigma-Aldrich). Five or six duplicates of each treatment were performed in each experiment.

### Golgi staining protocol

Golgi staining and spine counting were performed as described previously(*94*). Nine littermate pairs of male juvenile mice were used for this experiment. Golgi staining of the mouse brain was performed as instructed in the user manual for the FD Rapid GolgiStain™ Kit (FD NeuroTechnologies, Columbia, MD, USA). In brief, first, the mice were intracardially perfused with 0.9% saline. Next, the brain was dissected. The dissected brains were first immersed in the impregnation solution (solution A+B) for 21 days in the dark, then incubated in solution C for 3 days before slicing. A vibratome was used to cut 100 μm-thick coronal sections of mouse brains. After slide preparation, the sections were incubated in staining solutions (Solution D/E) for 10 min and washed twice with distilled water after staining. The samples were then dehydrated by sequential rinsing in 50%, 75%, 95%, and 100% ethanol, cleared with xylenes, and mounted with a mounting medium. A Nikon-TL confocal microscope was used to capture the images. To quantify the spine density, the images of one dendritic branch of each of four randomly selected somatosensory cortex neurons were taken from each sample. The density of dendritic spines was analyzed using Image J software.

### Confocal microscopy for mouse brain sections

Coronal sections of the cortical region (20 μm thick) were processed for dual immunofluorescence. In brief, the sections were incubated in blocking buffer, followed by mouse anti-NeuN (1:500), rabbit anti-NR1 (1:200), anti-mouse 3-Ntyr (1:200), and rabbit anti-GAD1 antibodies (each diluted 1:200). Then, the sections were rinsed with PBS and incubated with anti-rabbit Alexa fluor 594 (1:1000) and anti-mouse Alexa Fluor 488 (1:1000) secondary antibodies for 2 hours in the dark. After secondary incubation, the sections were washed with PBS 3 times and mounted on glass slides with DAPI. Images were captured at 100 x with a Nikon confocal microscope.

### Confocal microscopy for cultured cells

Cells cultured on coverslips were washed three times with PBS and fixed with 4% paraformaldehyde for 15 min at room temperature. The fixed cells were washed several times with PBS and incubated in the absence or presence of permeabilization buffer (PBS containing 0.1% Triton X-100) for 5 min at room temperature. After washing the cells three times with PBS, they were blocked with blocking buffer (PBS containing 5% BSA) for 30 min at room temperature and then incubated with the primary antibodies overnight at 4^0^C. Then the samples were incubated with anti-rabbit Alexa fluor 594 (1:1000) and anti-mouse (Alexa Fluor 488 (1:1000) secondary antibodies for 2 h, washed with PBS, and mounted with the nucleus fluorescent probe DAPI. Finally, the cells were observed (65X) under a Nikon confocal microscope.

### Human participants

All experimental protocols were approved by the Shaare Zedek Medical Center Institutional Review Board (Jerusalem, Israel; IRB# 0501-20-SZMC, original approval 02 December 2020). Whole blood samples were collected from 19 children with ASD and 20 unrelated, age- and gender-matched, neurotypical children who attended regular education classes and did not have any neuropsychiatric diagnosis. The demographic and symptom severity data of the cohorts are presented in Supplementary Table S2. Participants were recruited from the outpatient clinics and through advertisements posted in the surrounding community of Jerusalem, Israel. The study and its methods were performed in accordance with approved protocols and relevant guidelines and regulations. informed consent was obtained.

### Whole Blood Sample Collection

Venous whole blood was collected in the morning hours from non-fasted individuals. Whole blood samples were centrifuged within 15 minutes from the blood draw, and the plasma aliquots were frozen immediately at − 80 °C until use.

### Sample preparation for SNOTRAP

First, 9 mg of protein from blood plasma were taken. Lysis buffer was added to make the volume up to 1 ml and kept at 4° C for 10 min. Next, it was centrifugated at 13,300 rpm for 12 min at 4°C. The supernatant was collected, and 0.5 to 1% of SDS was added. Then, 9 mg of protein (supernatant) were washed with 50mM HEPES buffer 3 times at 5000xg for 30min at 10°C. The washed sample was transferred to a new low-binding eppendorf, and then 1.5mM (the final concentration) of SNOTRAP was added to the washed sample. SNOTRAP was synthesized as described previously (*32*, *55*). The SNOTRAP was incubated with the sample for 2h in the dark at room temperature. Then SNOTRAP-labeled samples were transferred to a 10 MWCO filter tube and washed 3 times with 50 mM ammonium bicarbonate (ABC) at 5000xg for 25 min at 10°C. In parallel, the streptavidin beads (200ul) were washed with ABC buffer (100 mM ABC, 150 mM NaCl, 0.05% SDS) 2 times and with ABC 50 mM 1 time at 600xg for 2 min at 4°C. Then, the washed samples and streptavidin were mixed and kept in a shaking mode for 2h at room temperature. After incubation, the samples were centrifuged at 600xg for 2 min at 4°C. Next, the supernatant was used as a non-SNO protein. Then, streptavidin beads having SNO-protein were washed 3 times with ABC buffer at 600xg for 2 min at 4°C and 3 times with 50 mM ABC for 2 min at 600xg and at 4°C. SNO-proteins were eluted from the beads with TCEP (10 mM), blocked with IAM, trypsinized, desalted, and sent to mass spectrometry analysis (*95*).

### nanoLC-MS/MS analysis

MS analysis was performed using a Q Exactive Plus mass spectrometer (Thermo Fisher Scientific, Waltham, MA USA) coupled online to a nanoflow UHPLC instrument, Ultimate 3000 Dionex (Thermo Fisher Scientific, Waltham, MA USA). Peptides (0.45 μg, as estimated by O.D.280 nm) were separated over a non-linear gradient (0 − 80% acetonitrile) and run at a flow rate of 0.3 μl/min on a reverse phase 25-cm-long C18 column (75 μm ID, 2 μm, 100Å, Thermo PepMapRSLC) for 120 min. The survey scans (380–2,000 m/z, target value 3E6 charges, maximum ion injection times 50 ms) were acquired and followed by higher energy collisional dissociation (HCD) based fragmentation (normalized collision energy, 25%). A resolution of 70,000 was used for the survey scans, and up to the 15 dynamically chosen most abundant precursor ions, with “a peptide preferred” profile were fragmented (the isolation window was 1.6 m/z). The MS/MS scans were acquired at a resolution of 17,500 (target value 1E5 charges, the maximum ion injection times were 120 ms). Dynamic exclusion was 60 sec. Data were acquired using Xcalibur software (Thermo Scientific). To avoid a carryover, the column was washed with 80% acetonitrile and 0.1% formic acid for 25 min between samples.

### MS data analysis

Mass spectra data were processed using the MaxQuant computational platform, version 2.0.3.0. Peak lists were searched against the translated coding sequences of the human proteome obtained from Uniprot. The search included cysteine carbamidomethylation as a fixed modification and oxidation of methionine as variable modifications; it allowed up to two miscleavages. The match between runs option was used. Peptides with a length of at least seven amino acids were considered, and the required FDR was set to 1% at the peptide and protein levels. Relative protein quantification in MaxQuant was performed using the label-free quantification (LFQ) algorithm.

### Behavioral tests

All behavioral tests were recorded using a video camera mounted above each platform and analyzed using a video tracking system, Ethovision XT 15 (Noldus Information technology BV).

### Open Field Test

The motor activity of mice was tested in an open field consisting of a white plastic arena (60 cm × 60 cm). In the first session (habituation), mice were placed individually in the center of the arena and allowed to move freely for 5 min. The next day, the mouse was placed in the same field. The total distance traveled (cm) was measured by the Ethovision XT system. After each test, the arena was cleaned with 70% alcohol solution (*96*).

### Object Recognition Test

To evaluate cognition, particularly recognition memory, we tested two object recognition tests or a novel object recognition test (NOR). This test utilizes the tendency of mice to explore novel stimuli (*97*). The test consisted of a familiarization session and a test session. At 24 h before these sessions began, the mice were allowed to explore the arena without objects for 5 min to habituate to their surroundings. During the first 5 min session (the first day), the mice were left to explore two identical objects that could be found at constant locations, 15 cm from the sidewalls, in the already familiar white plastic arena. Twenty-four hours later, the mice were introduced to the arena for a test session in which one of the familiar objects was replaced by a novel object. Exploration was defined as directing their nose to the object at a distance of ≤1 cm and/or touching the object with their nose. The time spent by the mouse exploring each object was recorded for 5 min. The arena and the objects were cleaned with 70% alcohol after each session.

### Elevated Plus Maze Test

The elevated plus-maze (*98*) consisted of four arms (30 x 5 cm): two open and two closed. This test was used to assess anxiety-related behavior in rodents. The platform was made of white plexiglass. The apparatus was elevated 45 cm above the floor. The test was initiated by placing the mouse on the central platform of the maze, facing one of the open arms, and letting it move freely. Each session lasted 10 min. The time spent in the closed and open arms was recorded.

### Three-Chambered Social Test

A three-chamber sociability test was used to evaluate the interest of the mice in engaging in social interaction with a stranger mouse. Generally, the mice prefer to spend more time with another mouse (sociability), and the mice tend to investigate a novel mouse more than a familiar mouse (social novelty). The social test apparatus consisted of a transparent acrylic box divided into three chambers (*99*). Two cylindrical wire cages were placed in chambers 1 and 3. On the first day, the test animal was introduced to the middle chamber for the sociability test and adjusted for 10 minutes. On the second day, an unfamiliar mouse was introduced into a wire cage in one of the side chambers, and the other side chamber remained empty. The time spent by the test mouse to explore the wire cage with an unfamiliar mouse inside was recorded for 10 minutes (the sociability test). On the third day, another stranger mouse was introduced into a wire cage in one of the side chambers, and the other side chamber had a familiar one. The time spent exploring the familiar mice and the unfamiliar mouse was recorded for 10 minutes (the social memory test).

### Statistics and Bioinformatics

The mean and Standard Error of the mean (SEM) were calculated for the western blots, immunofluorescence, spine density counting, and behavioral experiments. Student’s twotailed t-test was used for two-group comparisons. A one-way ANOVA test with the Tukey post hoc test was used for multiple comparisons. Statistical details and methods used in each experiment can be found in the figure legends. For Biological Processes (BP) analysis, we uploaded the lists of all SNO proteins into STRING, version 11.5. Protein-protein interaction of SNO-proteins was also done by STRING (http://string-db.org) (*100*). High confidence interactions (score > 0.7) from the neighborhood, gene fusion, co-occurrence, co-expression, experiments, databases, and text-mining lists were used. The quantification was based on the ion intensity of the peptides. We used Benjamini-Hochberg correction (*101*) on the p-value to generate a False Discovery Rate (FDR), and processes/terms with FDR values below 0.05 were included. Bar graphs and statistical analysis were performed using Prism 9.3 (GraphPad Software, San Diego, CA). p< 0.05 is considered statistically significant. p values are provided in the figure legends (*p < 0.05; **p < 0.01 ***p < 0.001).

## Acknowledgments

This work was funded by a US Department of Defense (DoD) grant, and an Israeli Science Foundation (ISF) grant, and a National Institute of Psychobiology in Israel (NIPI) grant, and an Israeli Council for Higher Education Maof Grant, and a Berettler Centre for Research in Molecular Pharmacology and Therapeutics Grant. We also thank the Satell Family Foundation and the Neubauer Family Foundation for their support.

