## Supplementary material for "The NO Answer for Autism Spectrum Disorder": Supp Info

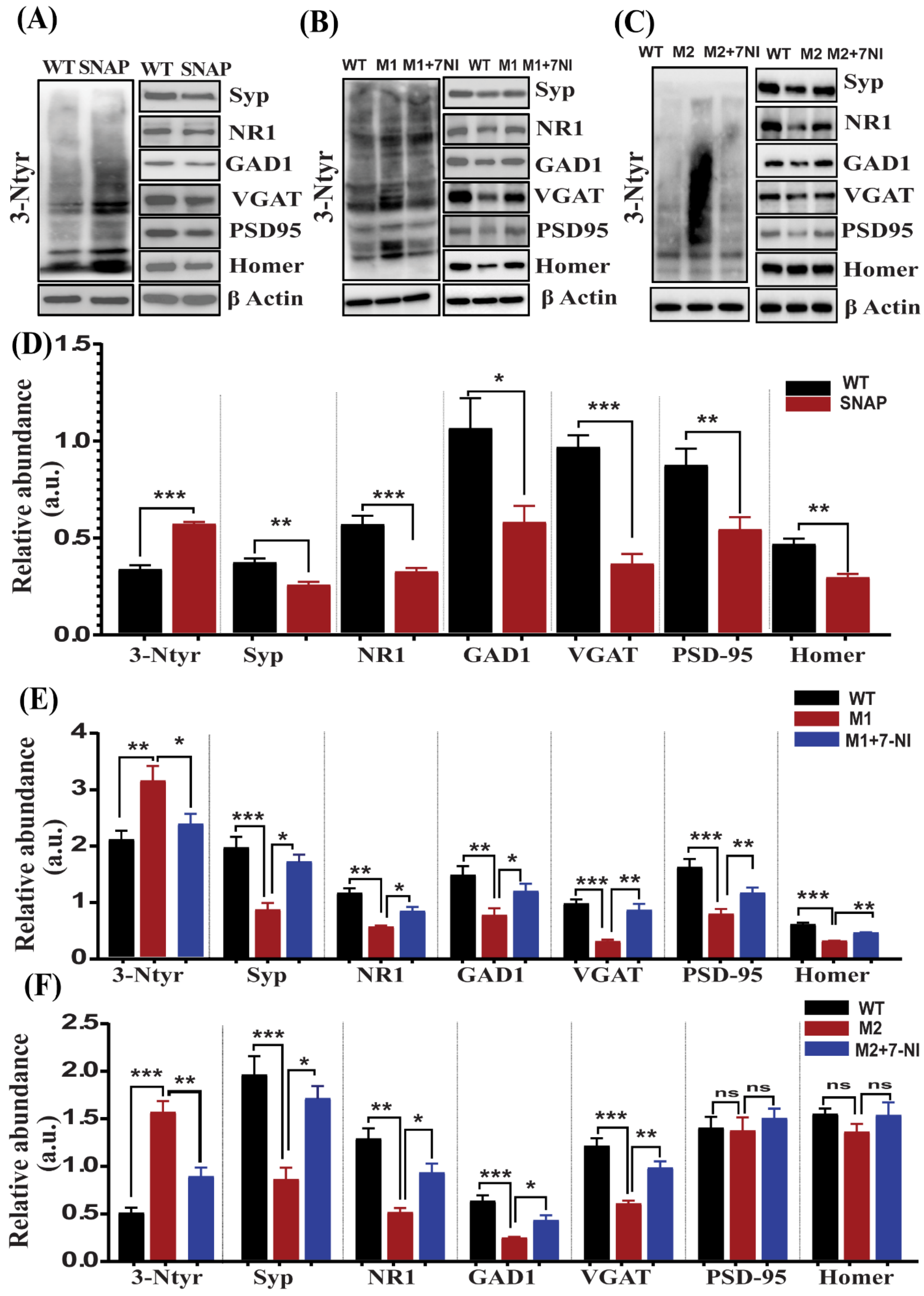

(A)
WT SNAP WT SNAP
3-Ntyr
Syp
NR1
GAD1
VGAT
PSD95
Homer
β Actin
(B)
WT M1 M1+7NI WT M1 M1+7NI
3-Ntyr
Syp
NR1
GAD1
VGAT
PSD95
Homer
β Actin
(C)
WT M2 M2+7NI WT M2 M2+7NI
3-Ntyr
Syp
NR1
GAD1
VGAT
PSD95
Homer
β Actin
(D)
Relative abundance (a.u.)
WT
SNAP
3-Ntyr Syp NR1 GAD1 VGAT PSD-95 Homer
(E)
Relative abundance (a.u.)
WT
M1
M1+7-NI
3-Ntyr Syp NR1 GAD1 VGAT PSD-95 Homer
(F)
Relative abundance (a.u.)
WT
M2
M2+7-NI
3-Ntyr Syp NR1 GAD1 VGAT PSD-95 Homer

**Figure S1. NO donor administration leads to nitrosative stress and synaptic pathology in the striatum of WT mice, and NO inhibition in *Shank3*$^{\Delta 4\text{-}22}$ mutant (M1), *Cntnap2*$^{(-/-)}$ mutant (M2) mice model led to a reversal in the molecular and synaptic deficits in the striatum**

**Panel A:** Representative Western blots of an indicator of nitrosative stress 3-Ntyr, and synaptic proteins Syp, NR1, GAD1, VGAT, PSD95, and Homer. β-actin was used as a reference for protein loading. **Panel B:** Representative Western blots of an indicator of nitrosative stress 3-Ntyr, and synaptic proteins Syp, NR1, GAD1, VGAT, PSD95, and Homer. β-actin was used as a reference for protein loading. . **Panel C:** Representative Western blots of an indicator of nitrosative stress 3-Ntyr, and synaptic proteins Syp, NR1, GAD1, VGAT, PSD95, and Homer. β-actin was used as a reference for protein loading. **Panel D:** Statistical analysis of the relative abundance of proteins shown in **Panel A.** **Panel E:** Statistical analysis of the relative abundance of proteins shown in **Panel B. Panel F:** Statistical analysis of the relative abundance of proteins shown in **Panel C.** A one-way ANOVA test with the Tukey post hoc test was used for multiple comparisons in **E,** and **F.** Student's two-tailed t-test was used for two-group comparisons in D * $P<0.05$, ** $P<0.01$, *** $P<0.001$. Groups of mice: WT (n=6); SNAP (WT mice treated with the NO donor compound SNAP, n=6), M1 (n=6); and M1+7-NI (M1 mice treated with the nNOS inhibitor 7-NI; n=6), M2 (n=6); and M2+7-NI (M2 mice treated with the nNOS inhibitor 7-NI; n=6).

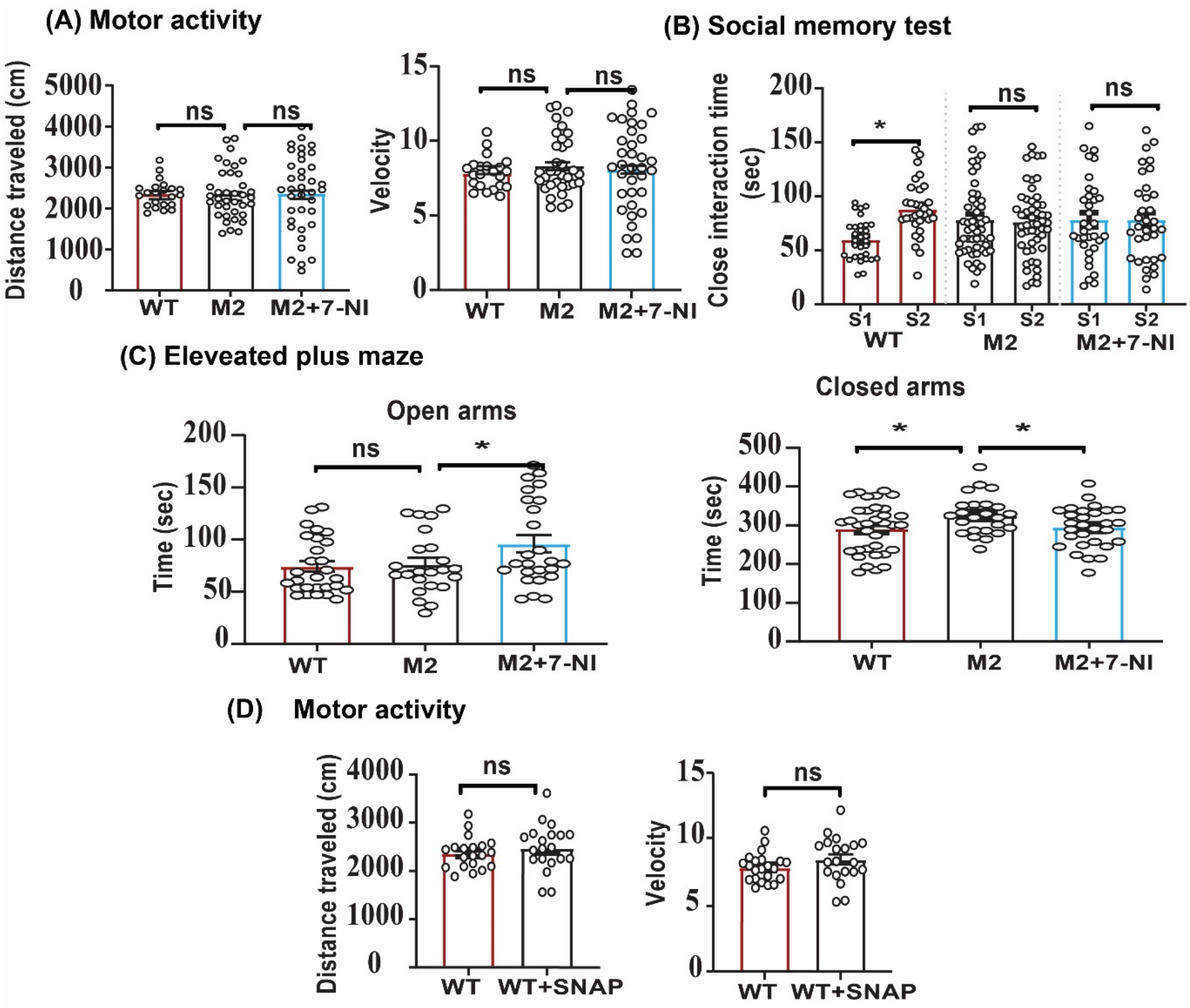

(A) Motor activity
Distance traveled (cm)
5000
4000
3000
2000
1000
0
ns
ns
WT
M2
M2+7-NI
Velocity
15
10
5
0
ns
ns
WT
M2
M2+7-NI
(B) Social memory test
Close interaction time (sec)
200
150
100
50
0
*
ns
ns
S1
S2
WT
S1
S2
M2
S1
S2
M2+7-NI
(C) Eleveated plus maze
Open arms
Time (sec)
200
150
100
50
0
ns
*
WT
M2
M2+7-NI
Closed arms
Time (sec)
500
400
300
200
100
0
*
*
WT
M2
M2+7-NI
(D) Motor activity
Distance traveled (cm)
4000
3000
2000
1000
0
ns
WT
WT+SNAP
Velocity
15
10
5
0
ns
WT
WT+SNAP

**Figure S2. 7NI-treatment corrects autistic behavior abnormalities in *Cntnap2*$^{(-/-)}$ mutant mice, and SNAP treatment increases behavioral abnormalities in WT mice**

**Panel A:** Statistical analysis of the motor activity in WT (n=22), M2 (n=37), and M2+7NI (n=37). **Panel B:** Statistical analysis of close interaction time with the novel intruder mice by WT (n=51), M2 (n=34), and M2+7NI (n=31) group. **Panel C:** Statistical analysis of total time spent in open arms and closed arms by WT (n=34), M2 (n=24), and M2+7NI (n=29). **Panel D:** Statistical analysis of motor activity test WT (n=22) and SNAP treatment group (n=20). A one-way ANOVA test with the Tukey post hoc test was used for multiple comparisons in A, B , C. Student's two-tailed t-test was used for two-group comparisons in D. * P<0.05, ** P<0.01, *** P<0.001, ns=not significant. Abbreviations, WT (control group), M2 (Cntnap2(-/-), M2+7-NI (Cntnap2(-/-) mice treated with the nNOS inhibitor 7-NI). SNAP (WT mice treated with the NO donor compound SNAP). S1=Familiar mouse, S2= novel mouse.

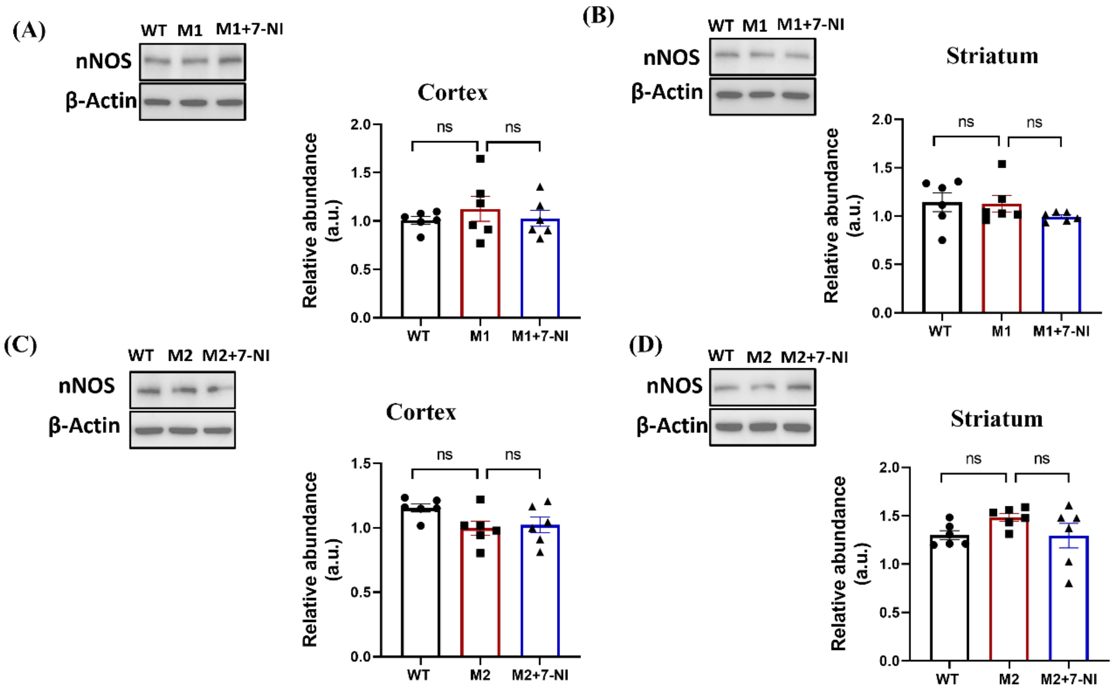


**Figure S3.** ***Shank3*$^{\Delta 4\text{-}22}$ mutation and *Cntnap2*$^{(-/-)}$ mutation do not change the protein expression of nNOS in mutant mice's cortex and striatum region.**

**Panel :** Representative western blots and statistical analysis of the relative abundance of proteins of nNOS in M1 group with and without 7-NI treatment in the cortex **(A)**, striatum **(B)**. Representative western blots and statistical analysis of the relative abundance of proteins of nNOS in theM2 group with and without 7-NI treatment in the cortex **(C)**, striatum **(D)**. Groups of mice: WT (n=6); M1 (n=6); M2(n=6), M1+7-NI (M1 mice treated with the nNOS inhibitor 7-NI; n=6), and M2+7-NI (M2 mice treated with the nNOS inhibitor 7-NI; n=6). A one-way ANOVA test with the Tukey post hoc test was used for multiple comparisons in all groups. * $P<0.05$, ** $P<0.01$, *** $P<0.001$, ns= not significant.

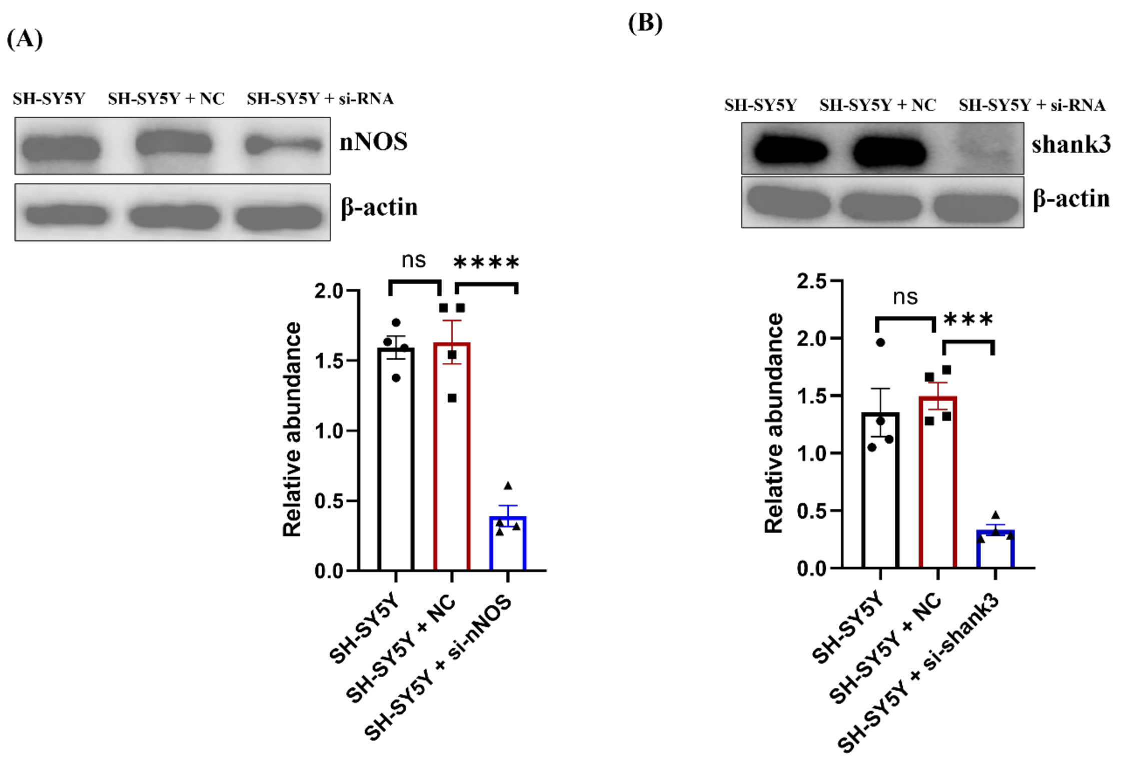


**Figure S4. Knockdown of *SHANK3* and *nNOS* expression in Human neuroblastoma cells by si-RNA.** Representative western blot of nNOS **(A)** and Shank3 **(B)** with statistical bar graph before and after siRNA treatment. A one-way ANOVA test with the Tukey post hoc test was used for multiple comparisons in all groups. * $P<0.05$, ** $P<0.01$, *** $P<0.001$, ns= not significant.

**Supplementary tables are uploaded as Excel/Word files:**

Table S1: MTT assay of 7-NI in primary cortical neuronal culture (Excel File).

Table S2: Clinical characteristics of participants (TD and ASD)

Table S3: IDs of SNO-proteins in the plasma sample of TD and ASD children (Excel File).

Table S4: System biology analysis of the plasma samples of TD and ASD children (Excel File).

**Table S2: Participant characteristics**

| | Neurotypical control | Children with ASD |
|---|---|---|
| ***N*** | 20 | 19 |
| Age | 4.6 | 4.2 |
| % male | 75% | 74% |
| **Symptoms severity** | | |
| ADOS Comparison score = 8-10 | | 63% |
| VABS Standard score ≤ 70 | | 84% |
| CARS Total score ≥ 37 | | 74% |
| SRS T-scores ≥ 75 | | 63% |
| MSEL, ELC ≤ 70 | | 63% |

Baseline characteristics of participants. There was no statistically significant difference between the group in age and sex. Only children with ASD had a neuropsychological evaluation. ADOS-2: Autism Diagnostic Observation Schedule, comparison score of 8-10 indicated severe autistic symptoms; CARS: Childhood Autism Rating Scale, scores above 36.5 are indicative of severe ASD; MSEL: Mullen Scales of Early Learning, ELC - early learning composite, scores below 70 are indicative of intellectual disability; SRS: Social Responsiveness Scale, Total Score ≥ 75 indicate severe autistic symptoms; VABS: Vineland Adaptive Behavior Scale, composite score ≤70 indicate low adaptive level.
